# Hidden Diversity of Threatened Sharks and Rays in the Global Meat Trade

**DOI:** 10.1101/2025.04.24.650194

**Authors:** M Aaron MacNeil, Chris G. Mull, Ana Barbosa Martins, Elizabeth A. Babcock, Zoya Tyabji, Alex Andorra, Shelley Clarke, Rima W Jabado, Glenn Sant, Joshua E. Cinner, Jessica A. Gephart, Nicholas K Dulvy, Arun Oakley-Cogan, Devanshi Kasana, Luke Warwick, Colin A. Simpfendorfer, Sarah Fowler, Marcio de Araújo Freire, Michel Bariche, Océane Beaufort, Joseph J. Bizzarro, Matias Braccini, João Bráullio de Luna Sales, Carlos Bustamante, John Carlson, Patricia Charvet, Juan Martín Cuevas, Cezar A. F. Fernandes, Daniel Fernando, Brittany Finucci, Emiliano Garcia Rodriguez, Adriana Gonzalez-Pestana, Luis Gustavo Cardoso, Rachel Ann Hauser-Davis, Efin Muttaqin, Maria del Pilar Blanco-Parra, Carlos J. Polo-Silva, Jonathan S Ready, David Ruiz-García, Luz Erandi Saldaña-Ruiz, Issah Seidu, Oscar Sosa-Nishizaki, Akshay Tanna, Rodolfo Vögler, Lucas Werner, Natascha Wosnick, Demian Chapman

## Abstract

International wildlife trade is a major source of biodiversity loss, yet many species lie hidden within aggregated data that conceals trade impacts. We overcome this problem for the largest vertebrate wildlife trade globally – shark and ray meat – comprising 438 538 mt yr^-1^ across more than 150 species, 76% of which are Threatened. Revealed trade contains greater quantities of skates (+10%), hammerheads (+8%), and smoothhounds, dogfishes & tope (+5%), and fewer pelagic sharks (-38%) than previously known. Shorttail yellownose skate, smoothound, silky, mako, and blue sharks are the most underreported meat species, due to aggregated landings from China, Argentina, Japan, and Indonesia, demonstrating international trade in shark and ray meat as a diverse, pervasive, and previously hidden source of fishing mortality for many threatened species.

---

International demand for shark and ray fins, combined with their high value and often non-perishable products (*1*), leads to long supply chains that hinder efforts to track source populations and develop effective conservation (*2*, *3*). However, beyond their fins, sharks and rays are fished for a wide range of other products – including meat, liver oil, gill plates, cartilage, and skins (*4–7*). Consequently, sharks and rays comprise the largest wildlife trade in the world in terms of number of animals killed and the likely diversity of species in unreported trade (*2*, *8*, *9*, *9–11*). While regulation of the fin trade has expanded to include 86% of traded species identified at major hubs (*12*, *13*), there has been no systematic exploration of the most fundamental driver of shark and ray exploitation – the market for meat.

Although trade flows in shark and ray meat are distinct (*6*), the species composition of the global meat trade remains largely unknown (*5*). Only one-third of shark and ray landings reported to the Food and Agriculture Organization of the United Nations (FAO) are identified to species (Supplemental Fig. 1), even where species-level information has been collected these may not be correctly identified and include complexes of congeners and confamilials. Two-thirds of landings are instead reported in “not elsewhere indicated” (NEI) categories that include an unknown number of species and trade is not reported to species, making it difficult to trace and detect the effects of meat trade on shark and ray populations. This lack of taxonomic resolution of landings has enabled overfishing resulting in steep catch declines and local extinctions (*14*).

Here we decompose the species composition of global trade in fresh or frozen shark and ray meat from 2012 to 2019, using integrated models of augmented FAO capture production data (see Appendix A) and reconciled Comtrade data from the Centre d’Études Prospectives et d’Informations Internationales (BACI; *Supplemental Information*). Our joint Bayesian model estimates the composition of landings available for trade from exporting nations using a combination of expert-elicited priors, reported and observed species-level landings, and a hierarchical decomposition of aggregated taxa (see Appendix B). This also allowed us to quantify information gaps in trade and landings data attributable to individual species and the countries catching them. We excluded analysis of re-exports, to ensure that we did not double-count volumes of shark meat in international trade. In essence, we combine the partially observed composition of species in reported landings with expert opinion to estimate the likely composition of species in the global meat trade.

We find that nearly half a million metric tonnes 438 538 [371 634, 535 472] mt (median [90% highest posterior density interval], live weight) of sharks and rays were traded annually for meat between 2012 to 2019 (Fig. 1A,B), representing some 120 [69, 205] million individuals each year (*Supplemental information*). Critically, although rays comprised one-quarter 29% [22%, 35%] of overall trade by live weight, due to smaller average body sizes they represent more than half 57% [40%, 77%] of individuals traded (Fig. 1C), reflecting the outsized and hidden impact of the global meat trade on often slower-growing and hence, more at-risk, ray populations globally (*15*). Shark meat was landed primarily by the longlining fleets of Spain, Taiwan, and Japan that dominate high-seas fisheries (Fig. 1D) and exported to Brazil and Europe (Fig. 1G), whereas trade in ray meat was dominated by exports of skate (*Rajidae*) from the United States and Argentina to South Korea and France (Fig. 1H).

**Figure 1.**
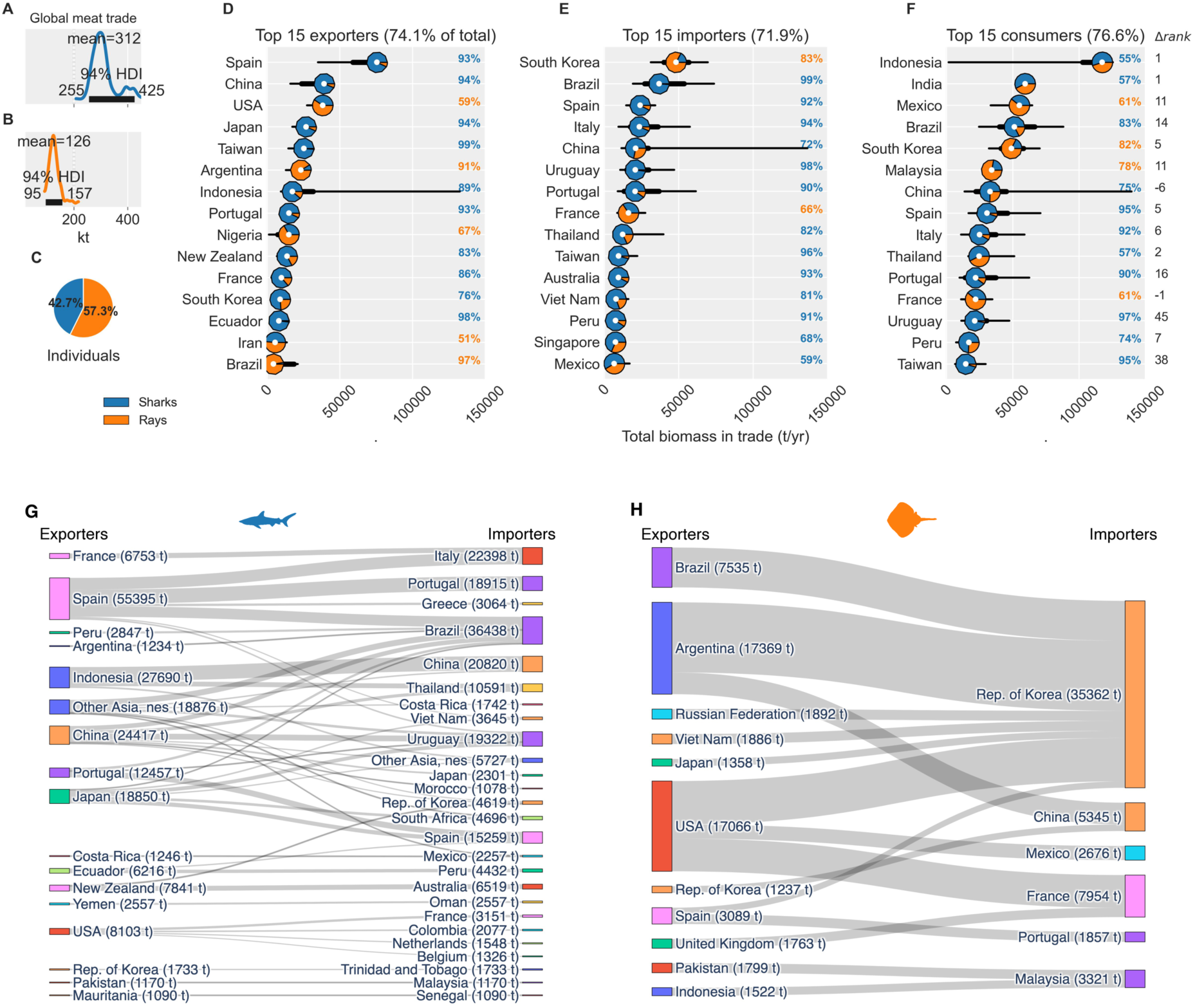
Estimated total weight (mt) of A) sharks and B) rays traded per year, with C) individuals; top 15 D) exporters, E) importers, and F) consumers (apparent consumption = retained landings + primary imports) in the global meat market for sharks and rays, 2012 to 2019; major (>1000 mt) global export links for G) sharks and H) rays, with estimated average tonnage in parentheses. Primary imports refers to biomass exported only once. White dots are median values, thick bars are 50% and thin bars 90% highest posterior density intervals. Proportions of sharks (blue) and rays (orange) indicated by donut plot colors, with dominant percentage listed at right in panels D-F. Difference between rank for apparent shark and ray meat consumption and rank for total seafood consumption (*Δrank*) listed at far right in F, with Spearman rank permutation correlation of 0.55 (p=0.0001) between ranked lists. Note difference in scale between G and H.

Making what is exported annually visible also reveals what is not – Indonesia, India, and Mexico retain most of their landings (Supplemental Fig 2) that, along with imports, make them the highest apparent consumers of shark and ray meat in the world (Fig. 1E). Yet compared to their global ranking of total seafood consumption, Indonesia and India do not preferentially consume shark and ray meat at the international scale (*Supplemental information*). Instead, we find that Mexico, South Korea, Brazil, Malaysia, Portugal, Spain, and Italy rank at least five positions higher in the retention and consumption of sharks and rays relative to other seafoods (*Δrank* in Fig. 1E; Spearman rank permutation correlation 0.55, p =0.001), suggesting strong cultural or economic demand for shark and ray meat sourced globally. These values include meat retained for other commodities, such as dried meat flowing from India to Sri Lanka and Nepal (*16*) and subsequent re-exports. For example, Uruguay also stands out as its shark and ray meat apparent consumption ranking (14^th^) is far higher than of general seafood (58^th^), due to its role as a major re-export hub for shark meat flowing mainly from Spain to Brazil in often unreported trade (*6*, *17*, *18*).

While blue shark (*Prionace glauca*) comprised 30% percent of annual global trade biomass, 156 other species make up the bulk of trade (>100mt/yr) – a diverse, but largely hidden, source of mortality for one-fifth of all shark and ray species (*Supplemental Table 2*). To characterize this diversity, we categorized all other shark and ray species within the major annual meat trade into eight priority fisheries groups that reflect distinct threats (*8*): pelagic sharks, hammerhead sharks, tropical requiem sharks, smoothhounds/dogfish/tope, skates, deepwater sharks, tropical stingrays, and ‘other’. Comparing the current state of species knowledge from reported FAO landings to our estimates for the global meat trade reveals a major shift in our understanding of species composition in trade, revealing greater contribution from skates (+9%), tropical sharks (+8%), and other coastal species (+8%), and away from pelagic sharks (-38%) than could be inferred from landings data alone (Fig. 2A). It is likely that higher species-level reporting within Regional Fisheries Management Organization (RFMO) landings that are dominated by pelagic species have downplayed the risks posed by the meat trade to non-pelagic species, which are more often hidden in aggregate, non-specified categories or absent from trade statistics altogether. The groups most traded for meat were skates, with 88% being exported primarily to South Korea and Europe, and smoothhounds/dogfish/tope, 78% of which were exported primarily to Europe and Australia.

**Figure 2.**
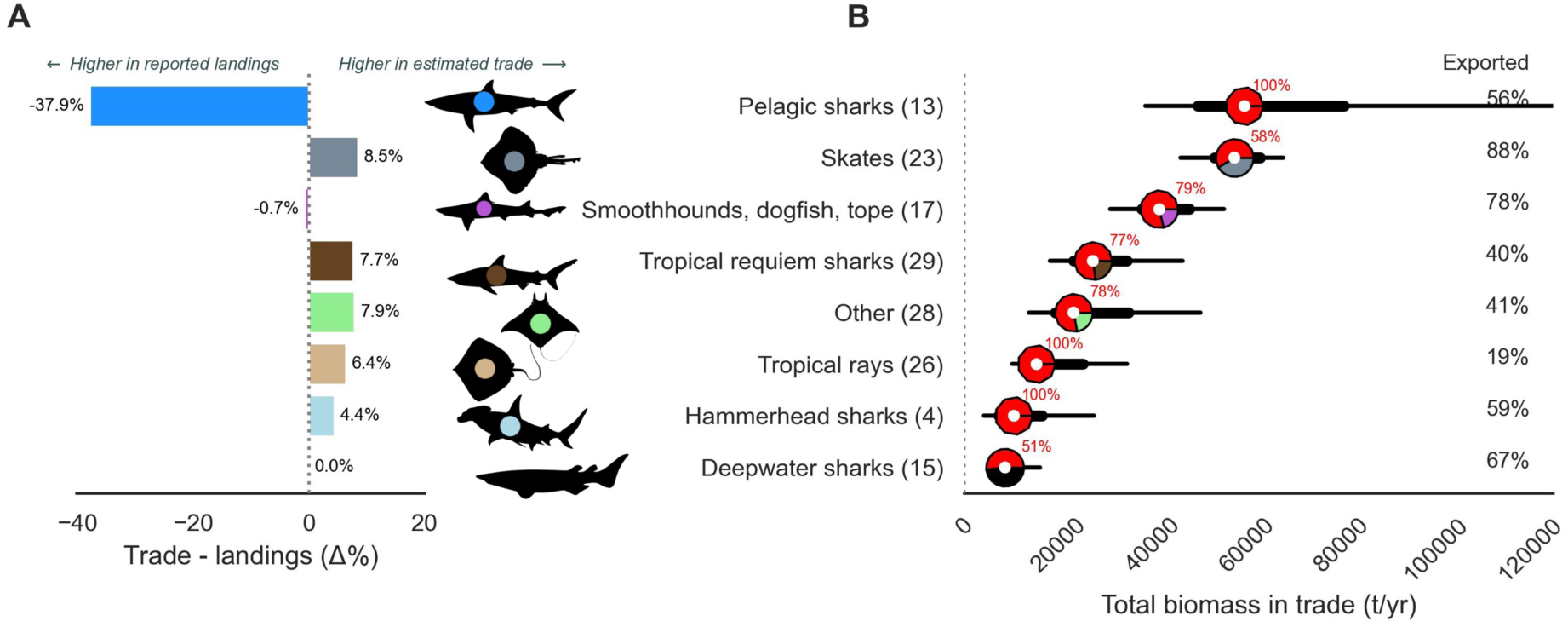
A) Estimated net differences in percentages of overall biomass (absent blue shark) among fished groups between species reported in FAO landings and species estimated within the global meat trade 2012 to 2019; B) estimated annual volume per fished group within the global meat trade 2012 to 2019. Numbers of species with >100 t of estimated trade for fished groups are given in parentheses; proportion of volume from threatened species (IUCN Red List status of ‘Vulnerable’ - VU, ‘Endangered’ - EN, or ‘Critically Endangered’ - CR c.2020) given in red; mean percentages of fished groups landed that are exported given in black at right. White dots in B) are median values, thick bars are 50% and thin bars 90% highest posterior density intervals. Colours match per fished group in A and B.

To understand the role of the international meat trade in contributing to species-level risk, we partitioned the 2012-2019 meat trade according to whether species were classified as threatened on the IUCN Red List of Threatened Species (*i.e.* Vulnerable, Endangered, or Critically Endangered) or otherwise. We find that 76% of non-blue shark meat traded from 2012 to 2019 came from species listed as threatened in 2020 (Fig. 2B), suggesting that the global meat trade has meaningfully contributed to declines in these species. This includes species highly susceptible to overexploitation with minimal to no management to halt fishing mortality, including deepwater species and tropical rays (*7*, *15*). We find that while the volume of skates in trade (∼57,000 t/yr) is similar to pelagic sharks excluding blue shark (∼59,000 t/yr), pelagic sharks have received far greater fisheries management and trade regulations attention (*19*). Skates are of increasing concern as they are often captured, and typically managed, as a species complex. For example, in Europe the four main skates are often landed together and reported as *Rajidae* or *Rajiformes* (20% of all European reporting) or may be binned as a single species (*e.g. R. clavata*). A key consequence of catch reporting and management of species aggregated is that it risks cryptic extinctions because the declines of larger species masked by increases in catch of smaller species (*20*, *21*).

The large percentage of threatened species in the global shark and ray meat trade raises the question as to how multilateral environmental agreements such as the Convention on International Trade in Endangered Species of Wild Fauna and Flora (CITES) and the Convention on Migratory Species (CMS), along with fisheries bodies such as tuna RFMOs, relate to the sharks and rays landed and traded for meat. To estimate this, we compared the 2012 CITES (Appendix I or II) and CMS (Appendix I or II) status, and RFMO measures of species landed in each country against their estimated volumes in both landings and trade using statistical models (see *Supplemental information*). Accepting at least 1–2-year implementation delays, we find that sharks and rays listed in such agreements continued to be landed and traded for meat – whether legally or illegally – throughout the study period, with a wide range of export proportions, from fully exported to almost entirely retained for local consumption (Fig. 3A). Four of the top five landed species overall – spiny dogfish (*Squalus acanthias*), shortfin mako (*Isurus oxyrinchus*), silky shark (*Carcharhinus falciformis*), and pelagic thresher (*Alopias pelagicus*) – are key targets for meat whose heavy exploitation led to their being listed under one or more measures (Fig. 3B).

**Figure 3.**
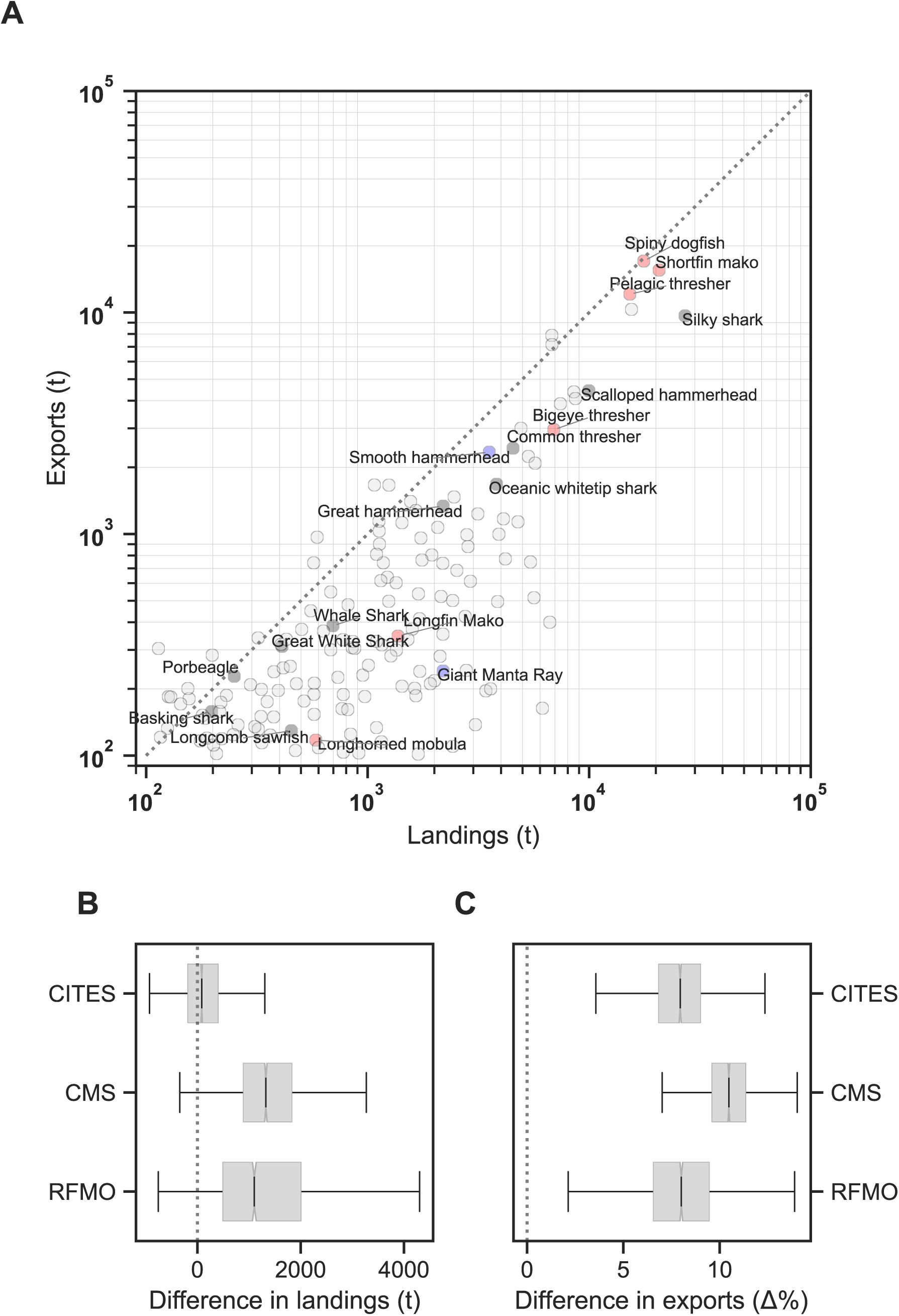
a) Estimated annual species exports versus landings volumes (>100 t) per country, 2012 to 2019; estimated effects of IUCN Red List threatened status (reflecting assessed low abundance) and CITES listing status on b) total landings and c) the proportion of landings exported per country and species in a). Colours in a) are species listed as IUCN Red List ‘Endangered’ or ‘Critically Endangered’ c2013 (red), CITES listing on Appendix II c2013 (blue), or both (black); grey points have none of these categories.

We find that, of the CITES species present in trade, only three – silky shark, shortfin mako, and scalloped hammerhead (*Sphyrna lewini*) – had more than 50 t/yr of meat trade reported only five times within the CITES trade database (Supplemental Table 1). By comparison, we estimate that an average 16901 [8314, 34546] t/yr of CITES listed species were landed, internationally traded, and not reported to CITES – demonstrating a wide gap between CITES reported trade and estimated trade shark or ray meat.

To quantify how listing under these trade and fishing agreements relate to international trade, we estimated their relationship to the percentage of landings exported per nation. Our estimates show that overall, CITES-listed species were 8% [5%, 11%] more likely to be traded on average than non-CITES species (Fig. 3C), regardless of landings volume (Fig 3B), while RFMO and CMS regulated species were equal or more likely to be exported than unlisted species, at 8% [4%, 12%] and 10% [8%, 13%] respectively. These findings show that international trade is a key contributor to overexploitation of species that merit listing and clearly demonstrates the magnitude of the CITES compliance gap for trade in shark and ray meat products, where the majority of CITES listed species in the international meat trade are not being reported.

There is a fundamental need to improve shark and ray data collection and reporting to the species-level (*22*) as countries with poor taxonomic resolution in catch and landings have been shown to have greater catch declines (*14*). The ongoing reporting of non-species-specific (i.e., NEI) catch and trade vastly underestimates fishing mortality and therefore the magnitude of threats to species hidden in landings and trade. While reporting of NEI categories within the FAO fisheries database declined slightly from 64% in 2012 to 62% in 2019, there has been no further improvement (*23*). Therefore, to quantify which species are underreported, and which countries contribute to underreporting in shark and ray landings and trade, we exploited a particular feature of our model: within each exporting country, we calculated a volume-weighted per-species information gap (𝐼𝐺_!_) based on Shannon’s information entropy. This metric reflects the probability of not being identified to species in the FAO database (see *Supplemental Information*) – a direct measure of the effects of aggregated reporting on understanding the species composition of landings that supply global meat trade.

We find that the most underreported species (landings by volume) occur in pelagic fisheries, where blue, shortfin mako, and silky sharks dominate estimated trade, along with thresher and hammerhead sharks (Fig. 4A). These CITES-listed species are also among those most prominent in global large-fin trade (*24*) and their estimated high volumes in the global meat trade demonstrates high use in multiple products that increases their risk of overfishing and severe depletion. While pelagic sharks have dominated international shark conservation advocacy and policy discussions for decades (*25*, *26*), our analysis reveals specific, meat-driven risks to several unlisted coastal species. These include shorttail yellownose skate (*Dipturus brevicaudatus,* VU) – taken extensively in coastal trawl fisheries on the Patagonian shelf and traded to South Korea. This species is not managed under any international agreement and species is not currently listed under CITES, likely representing a larger complex of undocumented species landings and trade flowing from Argentina to South Korea. Similarly, we find a large information gap for common smoothhound (*Mustelus mustelus,* EN) – a small shark that may also represent a larger species complex, exported for meat trade into France; smoothhounds and other *Triakidae* species have recently been found in high proportions of the Hong Kong small fin trade (*27*). Our analysis also reveals species for which information gaps are low – little skate (*Leucoraja erinacea*), winter skate, and tope (*Galeorhinus galeus*), which range from Least Concern to Critically Endangered on the IUCN Red List and all have high landings accurately reported by nations with strong governance and typically species-specific reporting practices such as the United States and New Zealand.

**Figure 4.**
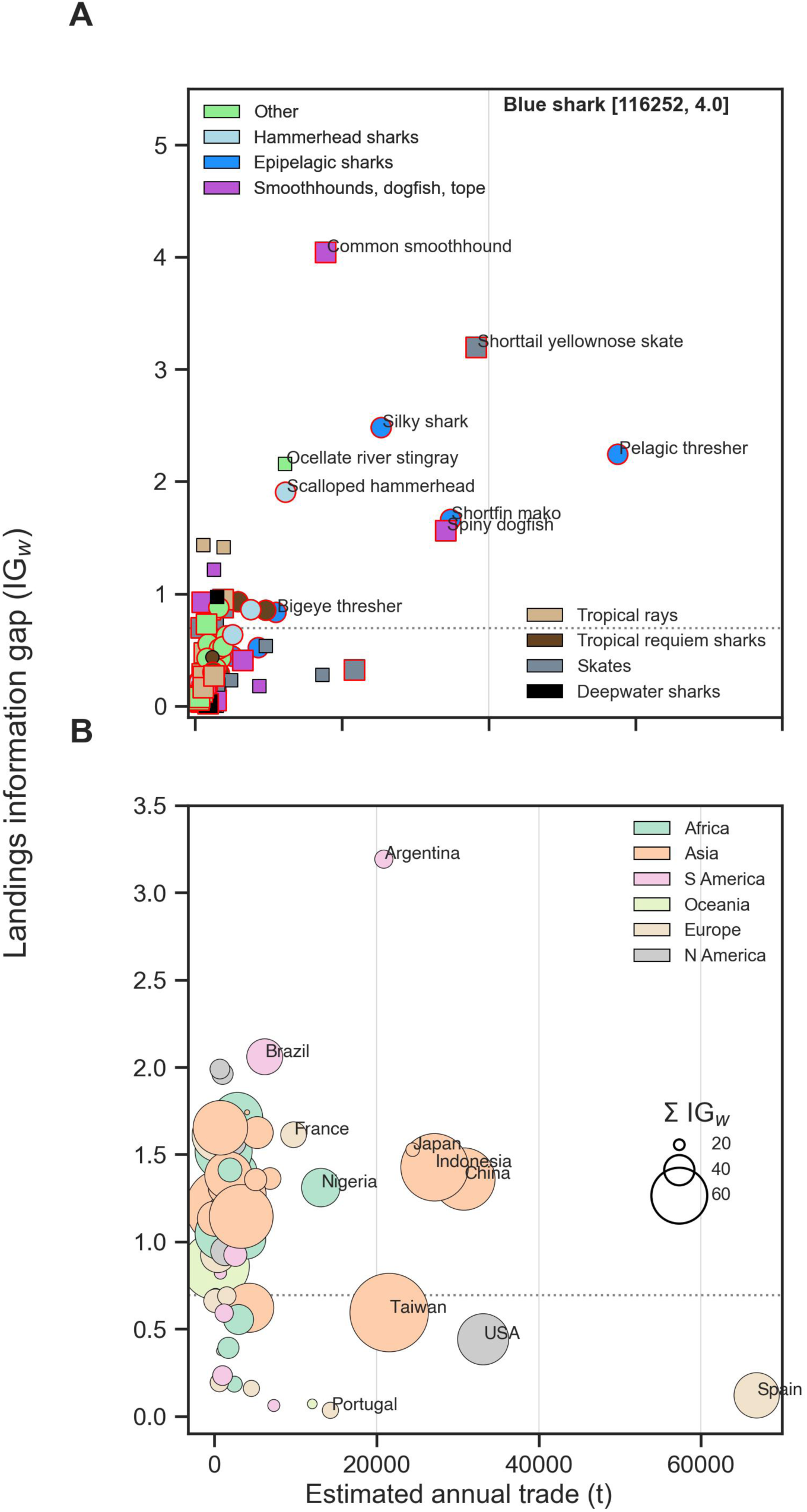
Information gaps across species and exporting nations relative to total trade in shark and ray meat excluding blue shark. a) Average landings information gap (in nats) per species weighted by their proportion of landings per country (red symbol edges are IUCN threatened species c.2020; circles are CITES listed species); b) information gap per country weighted by their proportional landings per species (circle sizes indicate total information gap across all species, Σ IG_w_). Information gap is the negative log-probability of being reported at the species level to FAO per country, called ‘self-information’ in information theory; horizontal dotted lines are equivalent to 50% reporting at the species level to FAO.

Global information gaps occur at the national scale, with countries choosing to report a wide range of taxonomic resolutions, from almost no NEI binning (*e.g*. Canada, New Zealand) to essentially no reported shark or ray landings of any kind (*e.g*. China, Vietnam, Thailand) – despite substantial estimated landings (Supplemental Fig. 4). Our analysis demonstrates that the largest information gaps for landings and trade in sharks and rays come from China, Argentina, Indonesia, and Japan – nations with high or diverse volumes of elasmobranch landings and substantial trade (Fig. 4B). These gaps occur even in cases where landings composition may be well documented nationally but are aggregated when reported to or by FAO. For example, Indonesia is the top apparent consumer of sharks and rays in the world, with a long cultural history of shark and ray meat consumption (*28–30*), with generally sufficient national level statistics to be used in non-detriment findings allowing trade of CITES-listed species (*31*). Yet of the ∼135,000 mt of sharks and rays landed by Indonesia annually, only three species were reported to FAO at the species level: blue shark (14,000 mt/yr), silky shark (5,800 mt/yr), and bottlenose wedgefish (*Rhynchobatus australiae*; 3,000 mt/yr) – were reported to FAO at the species level. And aside from 85mt/yr of Rajidae reported to FAO in 2012 and 2013, China reports no other shark or ray catches (FAO Fisheries and Aquaculture 2024, Pauly 2007, Zeller et al. 2016). Chinese fisheries expose a vast diversity of pelagic and coastal species to capture and the lack of species-level reporting contributes substantial uncertainty to the accuracy of global landings data. Many of the leading shark-fishing nations primarily operate artisanal or small-scale fleets, with landings often occurring at remote, unmonitored sites where reporting is absent. This lack of documentation likely leads to a significant underestimation of their contributions to both domestic and global trade, obscures the true composition of exploited, particularly coastal species, and hinders a comprehensive assessment of the ecological and socioeconomic consequences of shark and ray fisheries.

In contrast to those with high information gaps, other large exporters such as the USA, Spain, and Taiwan typically report most of their landings to the species level – in part due to RFMO reporting requirements – demonstrating that where data collection and reporting are highly resolved, information gaps are on average low. Importantly however, these fleets also represent 40% of total reported landings, and any unreported catch adds up quickly – while information gaps are on average low, absolute landings are large, resulting in large total gaps (Σ𝐼𝐺; Fig. 4B). Overall, the information gaps we identify are conservative in that they represent gaps only in reported landings – no bilateral meat trade reported at the species level and only a handful of unilateral meat trade is reported to FAO, amounting to less than 10 records 2012 to 2019.

Efforts to conserve shark and ray populations have gained momentum in recent decades, yet evidence suggests that adopted policies have yet to implemented with sufficient efficacy and scale to reverse increases in global shark landings and halt declines (*11*, *32*). It has the decades taken decades to establish the fragmented management measures within RFMO’s and to partially regulate regularly traded species via CITES Appendix II listings (*14*, *33*). Indeed, while finning regulations have reduced mortality at sea (Fowler and Seret 2010), they also increased landings of previously discarded carcasses. Furthermore, not all listings are legally binding, such as the International Plan Of Action (IPOA) sharks and associated National Plans Of Action (NPOA’s) and the CMS Sharks MOU (*14*). CITES Appendix II legality and sustainability certification requirements may limit exports and can result in associated changes to fisheries management practices, but do not directly affect domestic catch and consumption.

While addressing overfishing requires a range of polices across multiple dimensions from ocean to table, our results demonstrate that global meat trade is a key driver of exploitation that threatens a diverse range of shark and ray species. This information demonstrates that noncompliance with existing CITES listings is widespread within the meat trade. It also helps identify species caught without regulatory oversight that could benefit from international conservation and management measures given their prevalence in trade and creates the scope for a range of policy interventions, including implementing species-specific reporting through the supply chain, improved enforcement of species protections, bycatch reduction measures, better understanding of discard mortality, development of alternative livelihood options, monitoring and enforcement of CITES regulations in reporting meat trade, and national science-based catch quotas where possible. These measures, among others (*34*), will also ensure improved sustainability of high value and culturally significant foods (*35*). Thus, trade analysis represents a critical dimension for shark and ray conservation worldwide and has identified a clear example of large-scale wildlife trade driving an entire class of species toward functional extinction.

## Supplemental information

*Landings decomposition*

To estimate the species composition of global trade in shark and ray meat requires first estimating the species composition of global shark and ray landings. Here we relied on landings reported to any taxonomic level in the Food and Agriculture Organization of the United Nations (FAO) fisheries landings database (*23*). Individual countries report fisheries landings to FAO at varying taxonomic levels, meaning that to resolve species composition of global trade we first had to partially decompose the FAO landings to the species level. Note that it is generally assumed that FAO landings data improves on a 4-to-5-year time-lag, as countries refine reporting from previous years; therefore, we analyzed landings and trade from 2012-2019, the most recent years that are now considered stable.

Where FAO landings are reported to the species level, we had only to estimate the average landings over our 7-year time window (Appendix B). At higher levels of taxonomic aggregation (what we term the ‘taxon’ levels), reported landings can be comprised of increasing numbers of species that fall within each taxon group, all the way up to ‘Elasmobranch NEI’, the most general subclass grouping that includes both sharks and rays. This reporting structure meant that for each country and reporting level, there were combinations that were possible (present in country and could be within one of the reported taxon levels) and impossible (not within any of the reported taxon levels) that could be cancelled out of our calculations using a series of numerical masks. ‘Present in country’ was assigned if the species is present in either a country’s exclusive economic zone (EEZ) or in FAO Fishing Areas fished by each country’s fleet based on IUCN Extent of Occurrence maps. Using these criteria, we encoded a species mask of 1/0 values for each possible/impossible combination per country that was used in the model to define the possible species set in each reported taxon level per country.

While several countries report landings data to FAO – at least to the level of elasmobranchs NEI – many do not. For example, China and Vietnam provide little to no taxonomic resolution for many of their fisheries, each reporting over one million tons per year of unidentified marine fish that could potentially include elasmobranch landings. To include all the potentially important shark-fishing nations, we estimated the proportion of unidentified landings that could be elasmobranchs in each exporting nation using a separate statistical model (Appendix A). This model predicted latent total elasmobranch landings using a linear model with predictor variables including the fishing gears used, fraction of catch that is pelagic, and number of FAO areas fished, with the latent elasmobranch landings were constrained to be in the interval between the reported elasmobranch landings and the reported landings plus an estimated fraction of the unidentified landings. The unidentified elasmobranch catches used in our model were set equal to the median estimated latent elasmobranch landings minus the landings reported in elasmobranch categories.

Having a set of reported taxonomic levels and possible species combinations, we statistically estimated a latent (partially observed) species composition matrix of reported landings in each country (‘Latent landings’ in Appendix B Fig. 1), using elicited log-odds of species-specific landings within each nation from in-country experts, including project collaborators and members of the IUCN Species Survival Commission Shark Specialist Group (SSG). Categories included not retained (-4.5; log-odds scale, *LO_e)_*, rarely retained (-3.5), occasionally retained (-2.5), retained (-1), commonly retained (1), and large quantities retained (4). Note that while recommendations are available for eliciting the uncertainty in Dirichlet distributions (*36*), these approaches require experts to provide expected proportions along with upper and lower bounds - an exceptionally challenging task when considering 50 or more species. Instead, our *LO_e_* values were used as expected values in a ∼N(*LO_e_*, 3) constructed prior for each possible species landed by each country that was then passed through a *softmax* (multinomial-logit) function, expressing a high, common level of uncertainty across all possible species by exporter combinations. Methodological development in high-dimensional compositions would serve to reduce the uncertainty inherent in our relative quantity approach.

As part of our larger study, we conducted fieldwork in eight major coastal shark and ray fishing nations – Belize, Brazil, Mexico, Argentina, Uruguay, India, Sri Lanka, Indonesia, and Philippines – collecting field data from a range of small and larger scale ports (landing sites or fishing harbors) to quantify the species composition of shark and ray landings. While these studies occurred after 2019, they provide more detailed information about landings composition than our elicited prior weights alone. We also collated all available landings species composition data we could find from national fishing bodies, grey literature searches, and collaborations with researchers. To integrate all available information, we developed a landings sub-model where we used our elicited priors to estimate the posterior log-odds of the species composition in each nation, with information from observed data flowing back into our estimated species proportions per exporter and therefore contributing to our latent landings estimates (Appendix B Fig. 1).

*Trade decomposition*

With a decomposition of potential meat landings in place, we then decomposed species level landings as exports from each country into the global shark and ray meat trade. Trade data are, however, poorly resolved and over our study period from 2012-2019 and almost never to the species level; only a few rare reports of meat trade made to CITES provide any resolution. Furthermore, reported exports from one country to another often do not align with reported imports from another country. Rather than attempting to resolve these issues ourselves, we utilized the CEPII BACI trade database which resolves discrepancies between reported imports and exports among trading nations for each reported Harmonized System (HS) commodity code (https://www.cepii.fr/CEPII/en/bdd_modele/bdd_modele_item.asp?id=37).

For our purposes, the BACI data included key known meat commodity codes under HS2012 from 2012 to 2019 – fresh shark (030281, 030447, 030456, 030496); frozen shark (030381); fresh ray (030282, 030497, 030448, 030457); and frozen ray (030382) – that we used as the basis of our meat trade decomposition, summed as either shark or ray trade and scaled to round weight (the same scale as landings) using conversion factors (*Supplemental information*). One code – 030488, 1.7% of overall trade – included both sharks and rays and was binned as ‘shark’ because the bulk of 030488 observations (>80%) were from nations landing primarily shark. Note that this excludes meat traded as general fresh fish, frozen fish, or dried fish, all of which likely contain elasmobranchs (sharks and rays) to some degree; however, unlike landings, we could not readily decompose these categories into elasmobranch vs teleost groups, meaning our analysis is based on reported shark or ray meat trade only. In countries where total exports exceeded total landings (*e.g.* Singapore) we trimmed exports to a maximum of total reported shark or ray landings, meaning countries could only export a maximum of what they reported in landings (Supplemental Fig. 3) eliminating re-exports that could not have been fished locally. A key complication in the Comtrade/BACI data is that, despite a 2012 HS 030571 code for Fish; edible offal; shark fins, from 2012 to 2017 the frozen and fresh shark codes also include frozen fins, while the 2017 to 2019 data separate them. To remove the fins proportion from the pre-2017 data, we calculated the proportions of fresh (030292) and frozen (030392) fins in total fresh and frozen shark commodities in the post 2017 data (HS2017) for each country and adjusted the pre-2017 data by removing the average proportion of fins traded. Lastly, as shark and ray meat are traded as processed carcasses that do not represent the weight of an entire animal, we put trade on the same scale as reported landings using conversion factors given of 1.6 for sharks and 2 for rays to convert meat biomass volumes into total biomass based on FAO data (*37*).

While our augmented data accounts for total elasmobranch landings relative to total fish landings, meat trade data includes only high-value products traded fresh or frozen. Large volumes of elasmobranchs from 2012 to 2019 were also dried, salted, or smoked and likely exported as ‘dried fish’, in contexts such as *Glaucostegus* or *Rhinobatos* species in West Africa (RW Jabado, *unpublished*) and sharks moving from India to Sri Lanka (D Fernando, *pers. comm.*). Any dried meat and other uses such as pet food (*38*) would be binned into domestic retention in our results.

To decompose global trade in shark and ray meat required allocating our estimated landings per species into shark and ray exports reported for each nation. The simplest assumption would be that these exports are proportional to the species composition of each country’s landings, scaled up by total trade. However, we instead developed a linear model for species allocation that included a key adjustment – the potential (latent) species preferences of importers (‘Species affinity’ in Appendix B) that allowed for higher import volumes from countries exporting any given species. This factor essentially estimated the potential correlation in the species composition of exporting countries to a given importer, representing the tendency of nations to import more from countries that have species they desire. The linear model also included a trade friction coefficient, whereby nations with high seafood trade volumes would be more likely to also trade in shark and ray meat, and a dyad (pairwise) level effect to adjust for trade between any two countries relative to the global average trade. Lastly, while the species allocation model assigns potential landings by species from each exporter to a range of possible importers, not all species are used for meat – many will be diverted into pet food, fertilizer, aquaria, or other commodities. Therefore, to predict meat trade volumes among nations, we factored out (multiplied by zero) those species listed by the IUCN SSG as not being relevant for the meat trade (*i.e.* ‘Use and Trade’ in each listing).

*Numbers in trade*

While the biomass of species traded overall was the objective of our study, it raised the question as to what number of individual animals are landed each year to account for their biomass. To estimate total numbers would require knowing the size distribution of landed species, to at least an approximate degree. However, shark and ray fisheries vary widely in terms of what they land in selective gears, some being juvenile only fisheries, and some mature only fisheries. Many countries land immature animals of big species in the same group as adults of small species and make no distinction between them. The best approximation we could do was in general numbers, assuming a common relative size for all species that, while certainly incorrect, provides a first approximation of relative numbers of individuals traded per species. Therefore, to estimate the total number of individuals per species involved in the global meat trade, we divided the posterior estimated total volume traded for each species by the size at 75% maturity (*e.g.* Moore et al. 2012) as listed in Fishbase (fishbase.org), then summed across all species for each iteration of our model posteriors. This provided a distribution of estimated individuals in the trade (Supplemental Figure 5), assuming a fixed size that itself would be expected to vary locally depending on level of exploitation, size selectivity of fishing gears, and local availability of age classes to fisheries. In fisheries where large numbers of juveniles are landed – which may be often – these numbers would be much higher.

*Global seafood consumption rankings*

To understand the degree to which countries preferentially import and retain shark and ray meat relative to other seafood alternatives we ranked nations according to their global per-capita seafood consumption in 2021, based on data from Our World in Data (https://ourworldindata.org/grapher/fish-and-seafood-consumption-per-capita?tab=table). Countries deviating from their seafood ranks provide evidence of preference or aversion to elasmobranch meat consumption relative to their overall seafood consumption.

*Model validation*

To check our trade composition against known information, we compared our estimates for species composition of the shark meat trade to those derived from genetic identification of shark fin clippings in the Hong Kong (HK) and Guangzu (GZ) fin markets (*24*), given as proportions of overall trade. Our estimates of species proportions within the meat trade closely matched those reported independently reported fin trade (Supplemental Figure 6), with HK and GZ relative proportions frequently being with the 50^th^ percentile of our posterior estimates, indicating little difference in proportions of species within either trade. One key exception was spiny dogfish (*Squalus acanthias*) a species specifically targeted for meat for using in fish and chips around the world – while dogfish were less than 0.2% of fin markets, they were some 4% [2%, 6%] of global meat trade, exceeding 14,000t annually. However their fins are traded into the parallel small fins trade (*27*, *40*).

*Information gaps*

To quantify where uncertainty lies in understanding the species composition of elasmobranch landings and trade, we calculated a volume-weighted per-species information gap (𝐼𝐺_!_) using the average negative log-probability of being identified to species in the FAO database (Supplemental Figure A1):

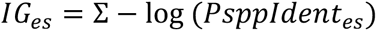

where *PsppIdent_es_* is the posterior distribution for the proportion of estimated latent landings that is estimated to be reported to FAO at the species level each year on average. It is a direct measure of the effects of aggregated reporting on the information available concerning the species composition of landings that supply global meat trade. Where the bulk of estimated landings are reported to the species level (*e.g.* Tope in New Zealand, *Spp ID logOdds* ∼0.97) the information gap is low (∼0.03), indicating there is little uncertainty in those reported data. Conversely, where the bulk species landings are not reported (*e.g.* blue shark in China ∼0.02), the information gap is high (∼3.9). While this decomposition was made on information gaps in reported landings to species rather than trade, because trade is unreported to species these estimates also represent the information gaps in trade – essentially, global ignorance of the meat trade is uniformly bad, making the two areas of ignorance directly proportional.

**Supplemental Table 1.** Trade in shark and ray meat reported to CITES 2012 to 2019. Data downloaded from CITES Trade Database (https://trade.cites.org/) October 2024.

| Year | App. | Taxon | Importer | Exporter | Importer reported quantity (mt) | Exporter reported quantity (mt) | Term |
| --- | --- | --- | --- | --- | --- | --- | --- |
| 2019 | II | Carcharhinus falciformis | MEX | CRI | 0 | 391 | meat |
| 2019 | II | Isurus oxyrinchus | KOR | ZAF | 0 | 141 | meat |
| 2016 | II | Sphyrna lewini | KOR | CHN | 0 | 95 | meat |
| 2017 | II | Sphyrna lewini | KOR | CHN | 0 | 83 | meat |
| 2019 | II | Isurus oxyrinchus | TWN | JPN | 0 | 50 | meat |
| 2019 | II | Carcharhinus falciformis | GTM | CRI | 0 | 46 | meat |
| 2019 | II | Isurus oxyrinchus | TWN | VUT | 0 | 35 | meat |
| 2019 | II | Carcharhinus falciformis | GTM | CRI | 25 | 0 | meat |
| 2014 | III | Lamna nasus | ESP | JPN | 24 | 0 | meat |
| 2019 | II | Carcharhinus falciformis | TW | CRI | 0 | 23 | meat |
| 2013 | III | Lamna nasus | ESP | JPN | 23 | 0 | meat |
| 2018 | II | Sphyrna lewini | KOR | CHN | 22 | 22 | meat |
| 2019 | II | Alopias pelagicus | USA | ECU | 0 | 15 | meat |
| 2018 | II | Carcharhinus falciformis | ESP | OMN | 0 | 9 | meat |
| 2019 | II | Carcharhinus falciformis | ITA | OMN | 0 | 5 | meat |
| 2019 | II | Isurus oxyrinchus | HKG | ESP | 0 | 23 | meat |
| 2014 | II | Lamna nasus | DNK | NOR | 1 | 1 | meat |
| 2015 | II | Lamna nasus | DK | NO | 1 | 1 | meat |
| 2014 | II | Mobula spp. | US | MX | 1 | 0 | meat |
| 2018 | II | Alopias vulpinus | US | CR | 1 | 0 | meat |
| 2014 | II | Mobula spp. | US | MX | 1 | 0 | meat |
| 2018 | II | Alopias pelagicus | US | CR | 0.2 | 0 | meat |
| 2016 | II | Lamna nasus | DK | NO | 0.1 | 0.1 | meat |
| 2015 | II | Lamna nasus | CN | CA | 0.1 | 0 | meat |
| 2015 | II | Lamna nasus | DK | FO | 0.1 | 0 | meat |
| 2015 | II | Lamna nasus | US | CA | 0 | 0.1 | meat |
| 2019 | II | Isurus oxyrinchus | US | LK | 0.1 | 0 | meat |
| 2015 | II | Mobula spp. | US | MX | 0.05 | 0 | meat |
| 2016 | II | Mobula spp. | US | MX | 0.03 | 0 | meat |
| 2016 | II | Mobula spp. | US | MX | 0.01 | 0 | meat |
| 2017 | II | Rhincodon typus | US | MX | 0.01 | 0 | meat |
| 2016 | II | Mobula spp. | US | MX | 0.01 | 0 | meat |
| 2017 | II | Mobula spp. | US | MX | 0.01 | 0 | meat |
| 2015 | II | Mobula spp. | US | MX | 0.001 | 0 | meat |

**Supplemental Table 2.**
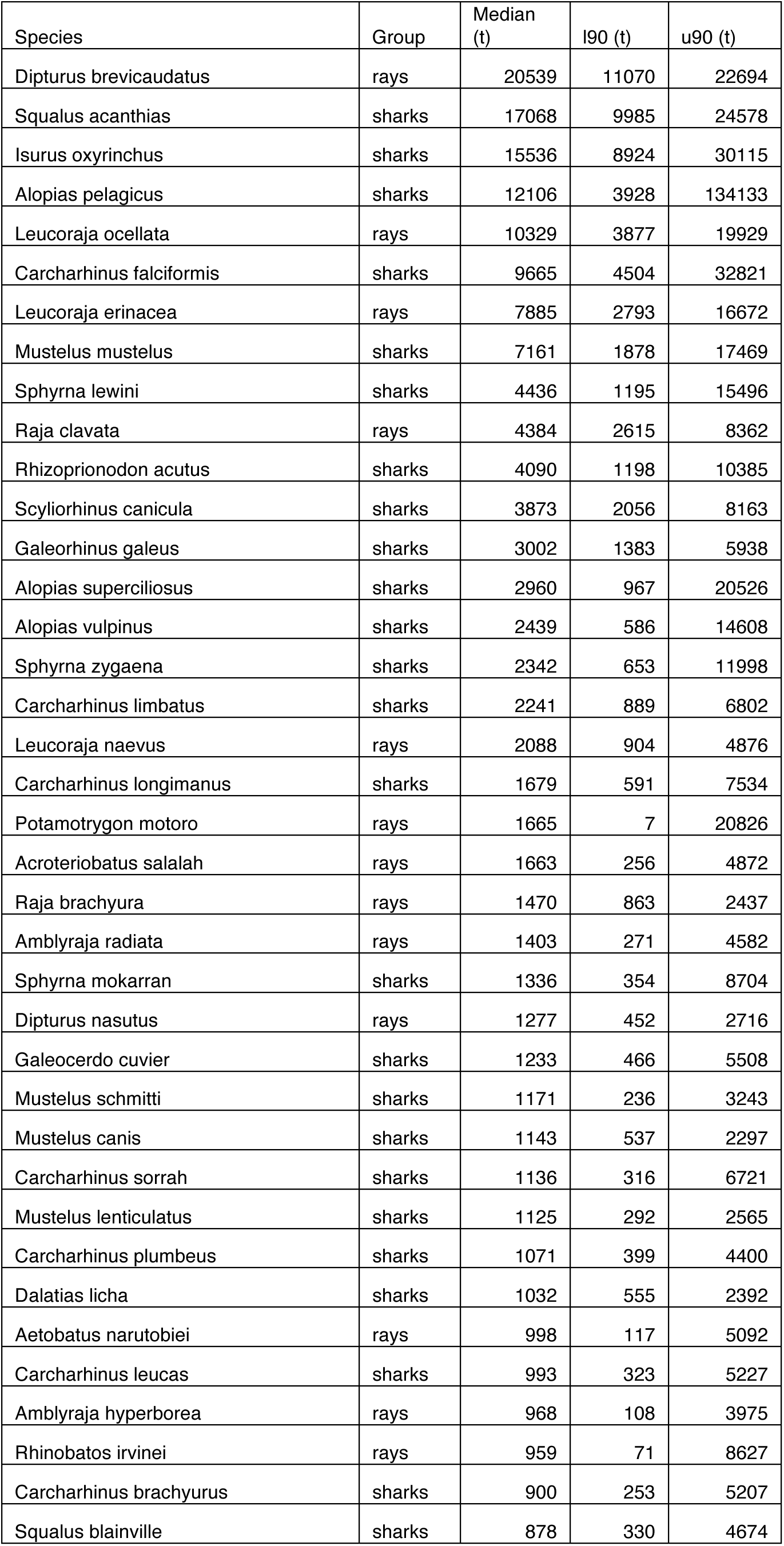
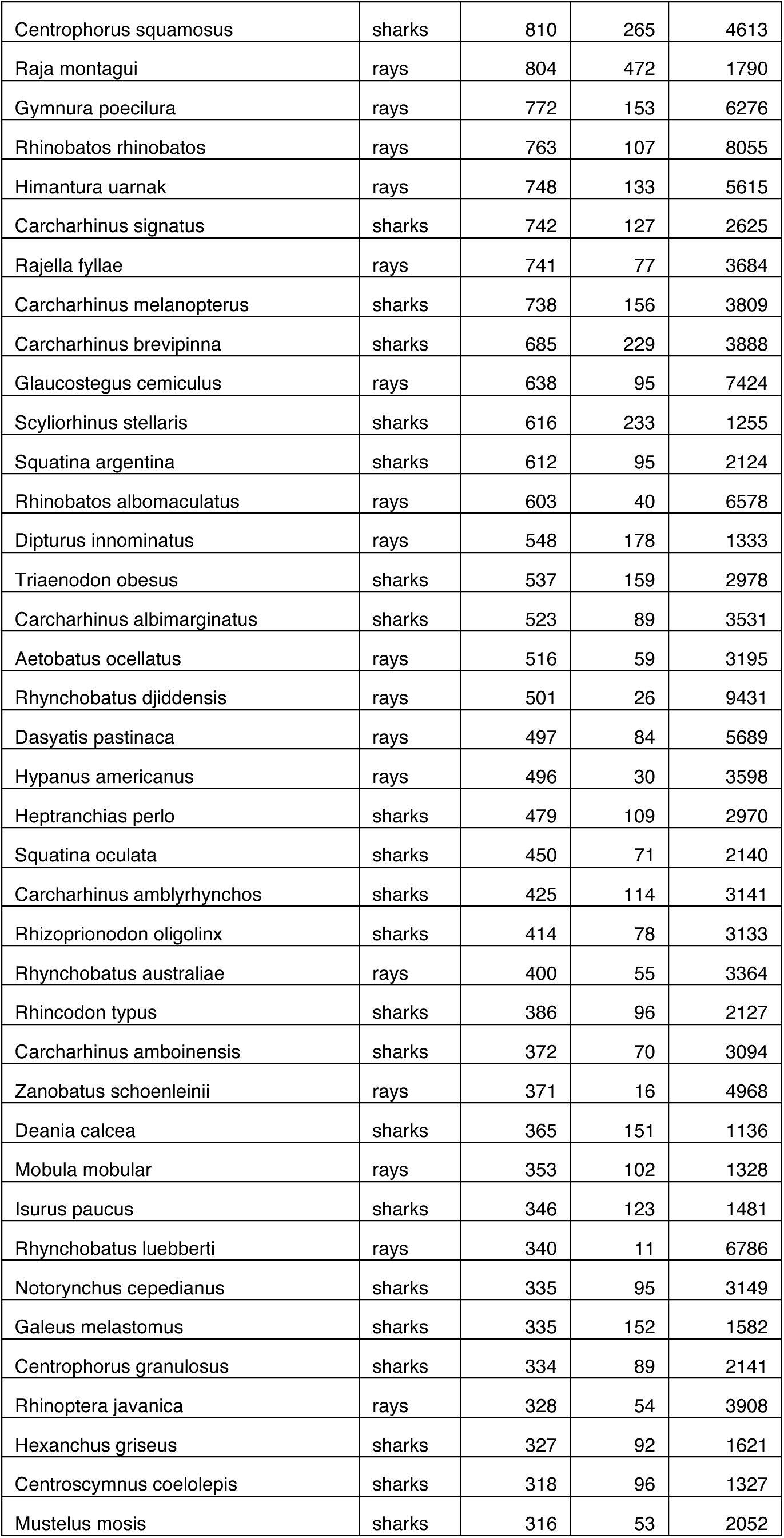
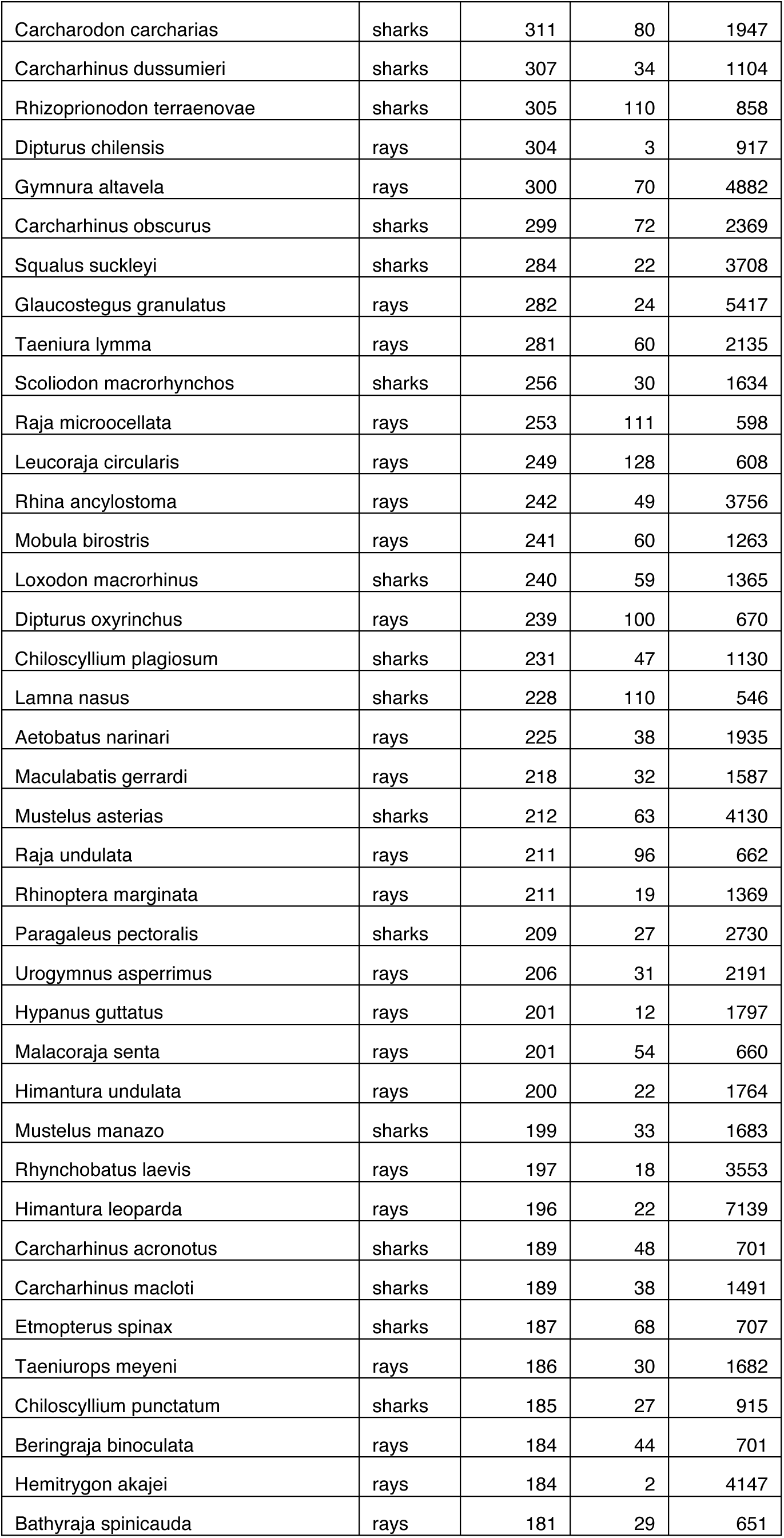
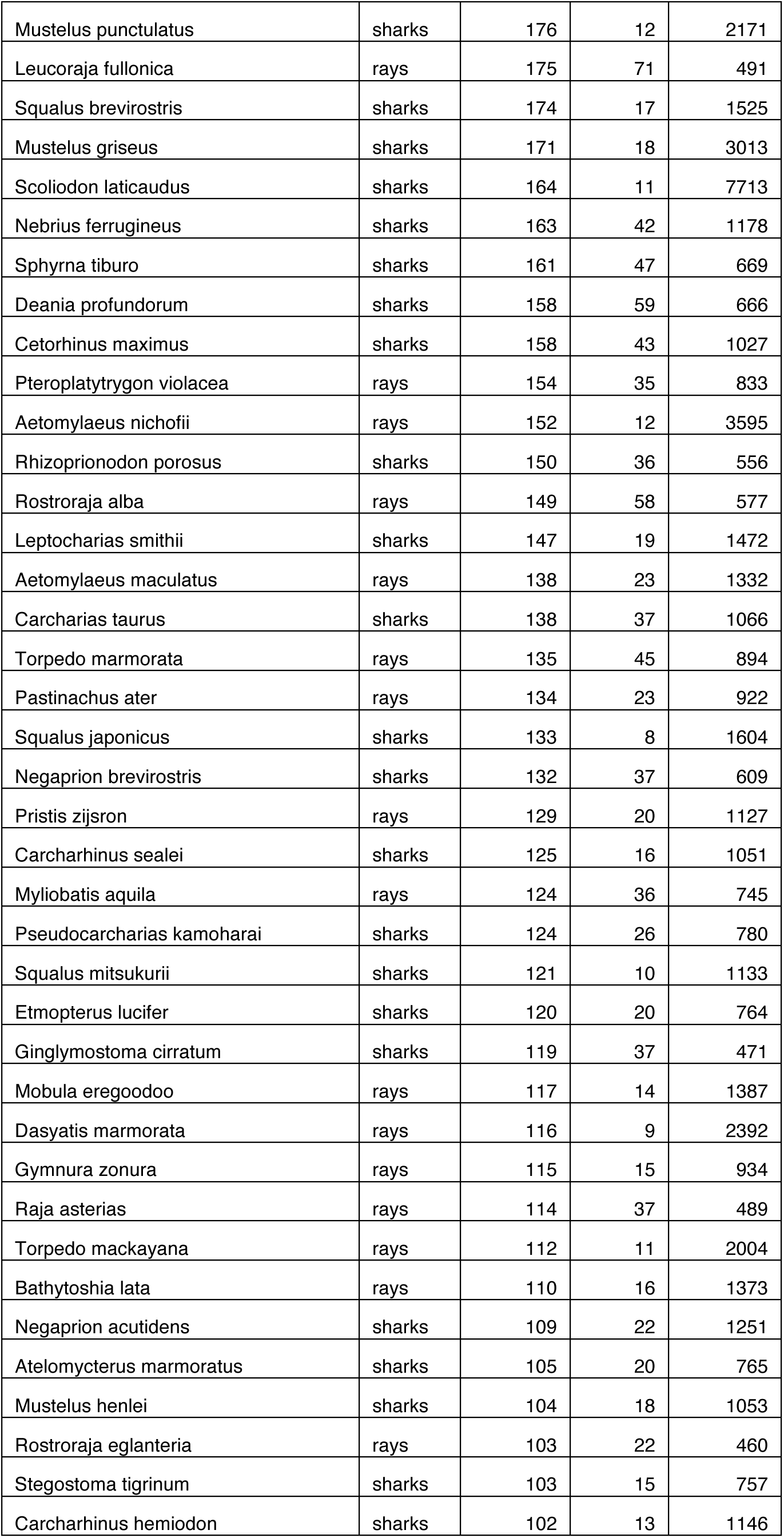
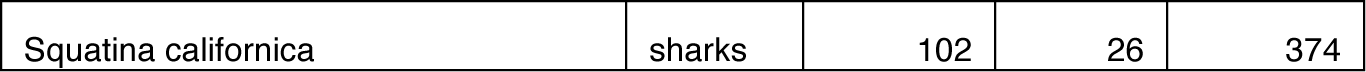
Estimated species traded for >100t/yr of meat, including medians, lower 90% (l90), and upper 90% (u90) highest posterior densities.

## Supplemental Figures

**Supplemental Figure 1.**
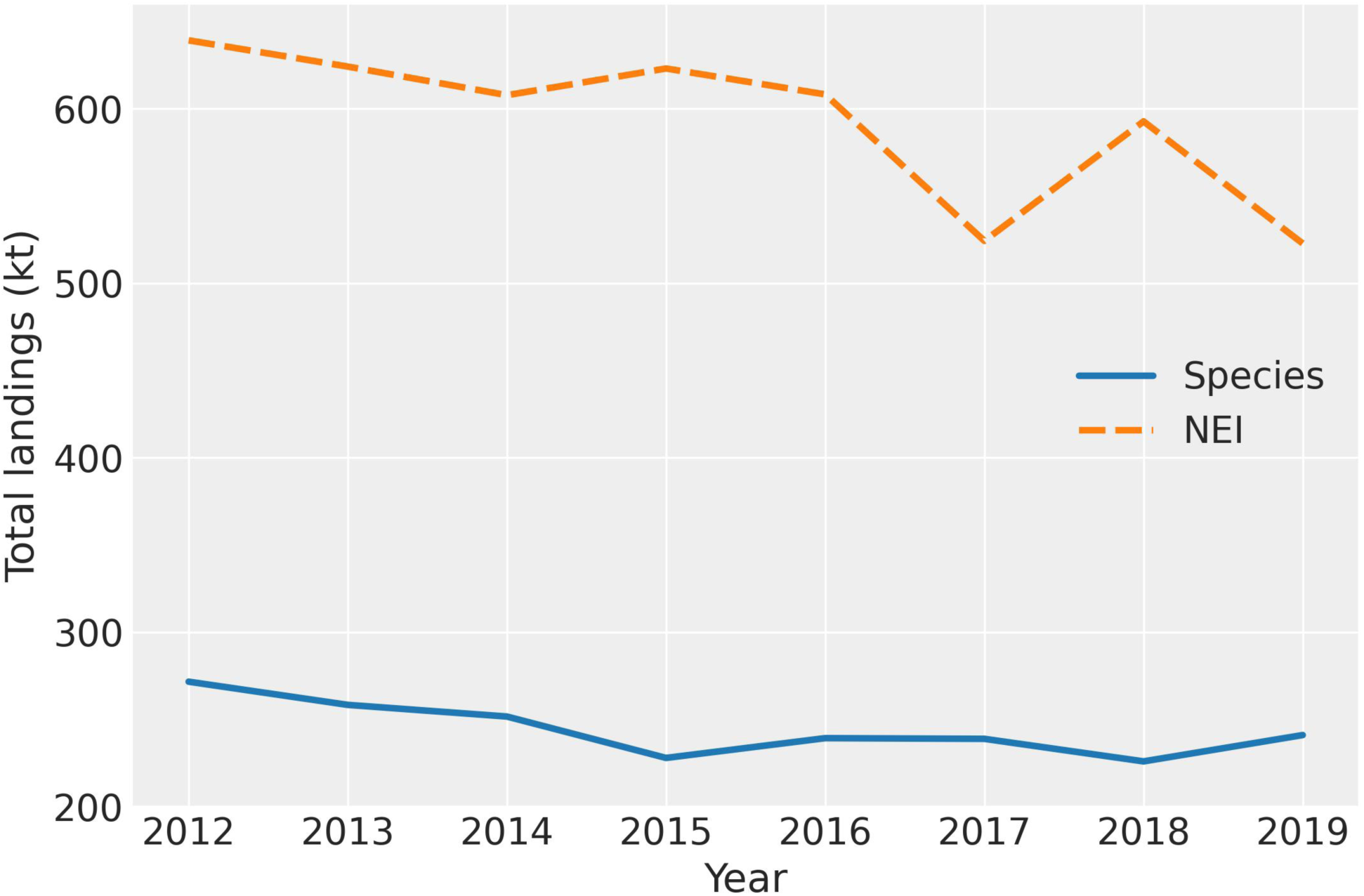
Total shark and ray landings reported to FAO at the species level (blue solid line) and some level of not elsewhere identified (NEI; orange dashed line) aggregation, 2012 to 2019.

**Supplemental Figure 2.**
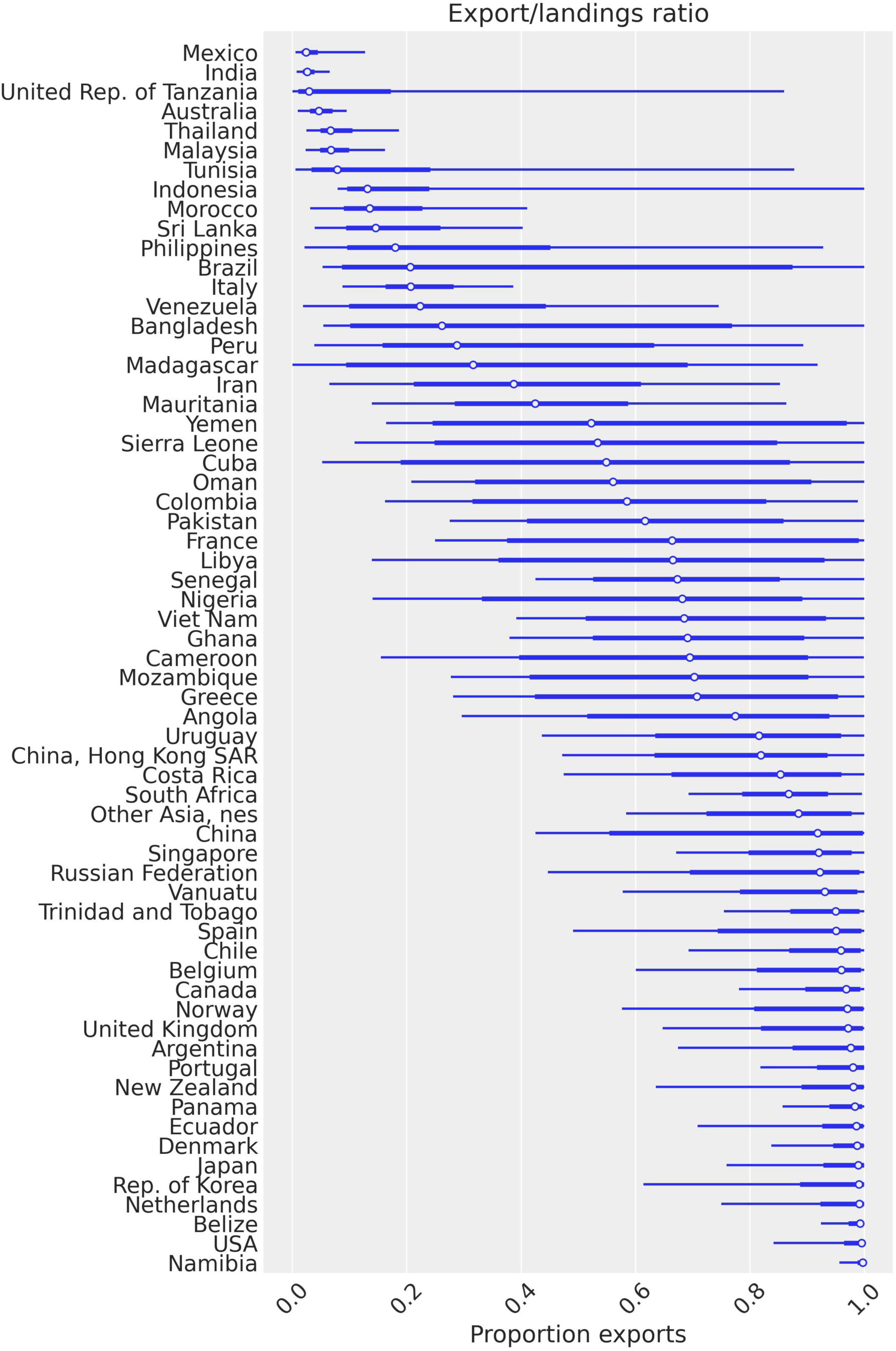
Estimated proportion of total reported landings exported per nation. Values of 1 indicate all reported landings are exported; trade values have had minimum re-exports removed (see Supplemental Figure 3).

**Supplemental Figure 3.**
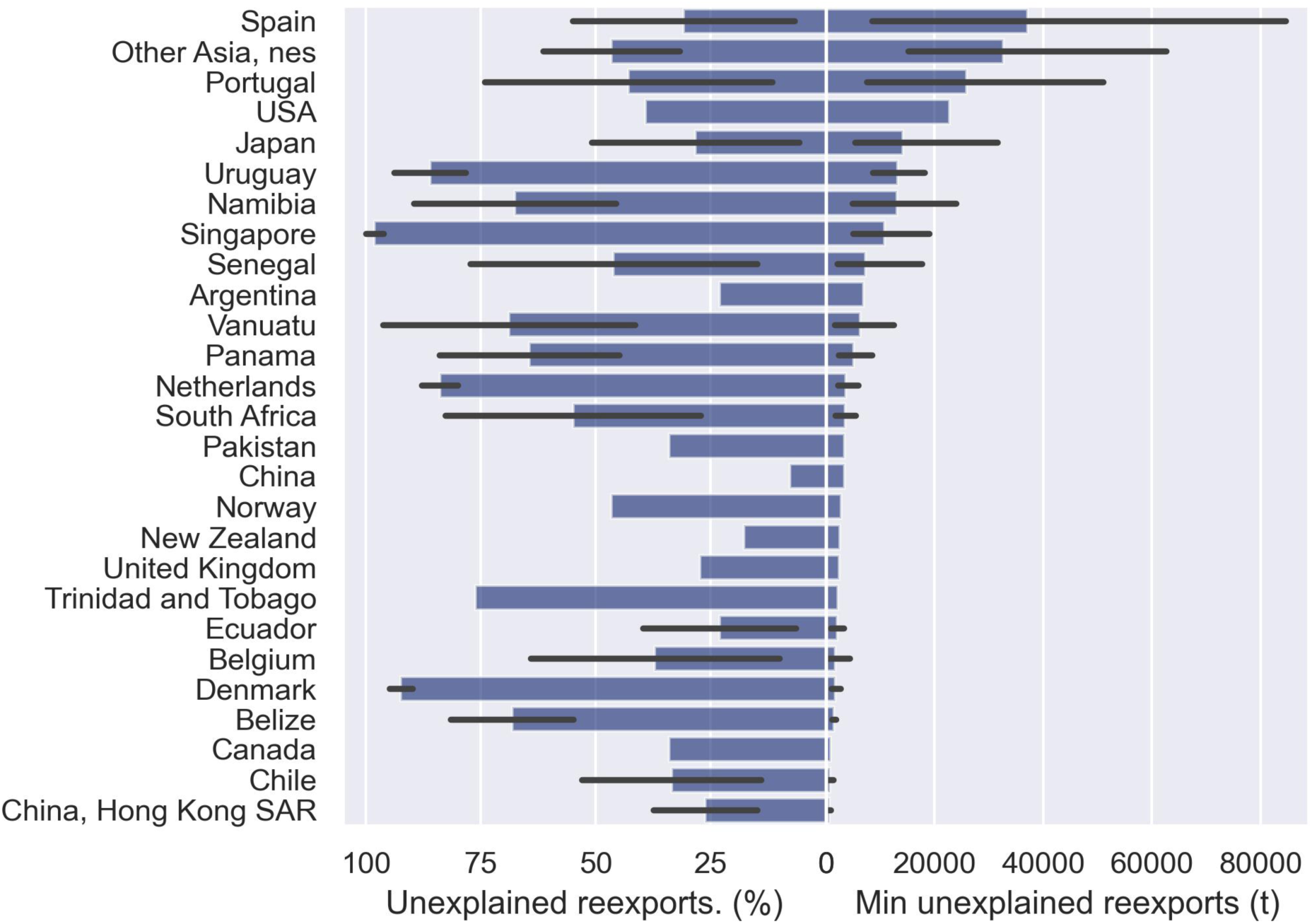
Percent and total minimum reexports for estimated landings and trade of shark and ray meat, 2012 to 2019. Values represent differences by which total BACI exports exceed augmented FAO landings, representing minimum reexports per year.

**Supplemental Figure 4.**
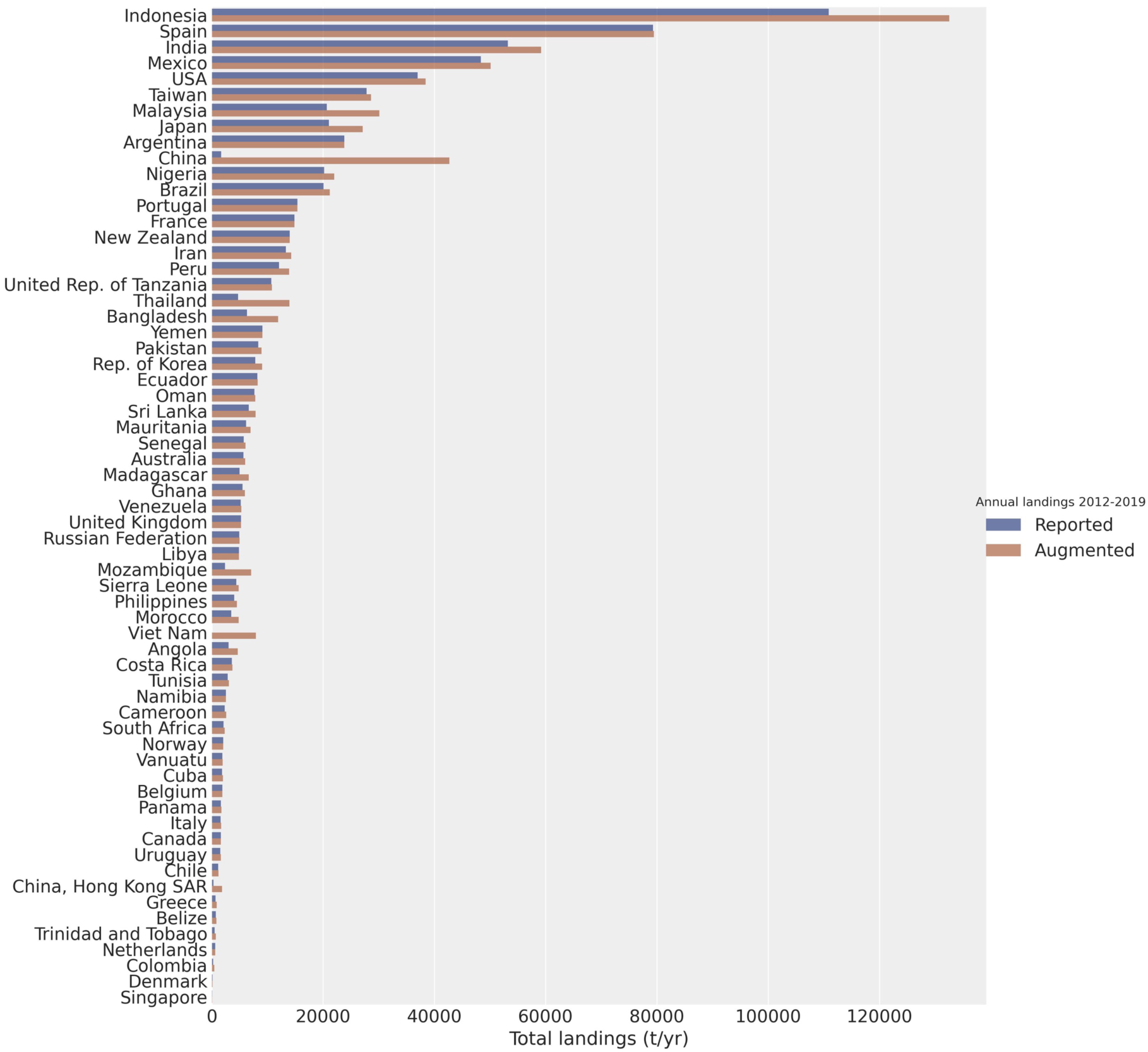
Reported and model-augmented average total shark and ray landings per year, 2012 to 2019. Discrepancies between barplots indicate countries estimated to be underreporting elasmobranch landings.

**Supplemental Figure 5.**
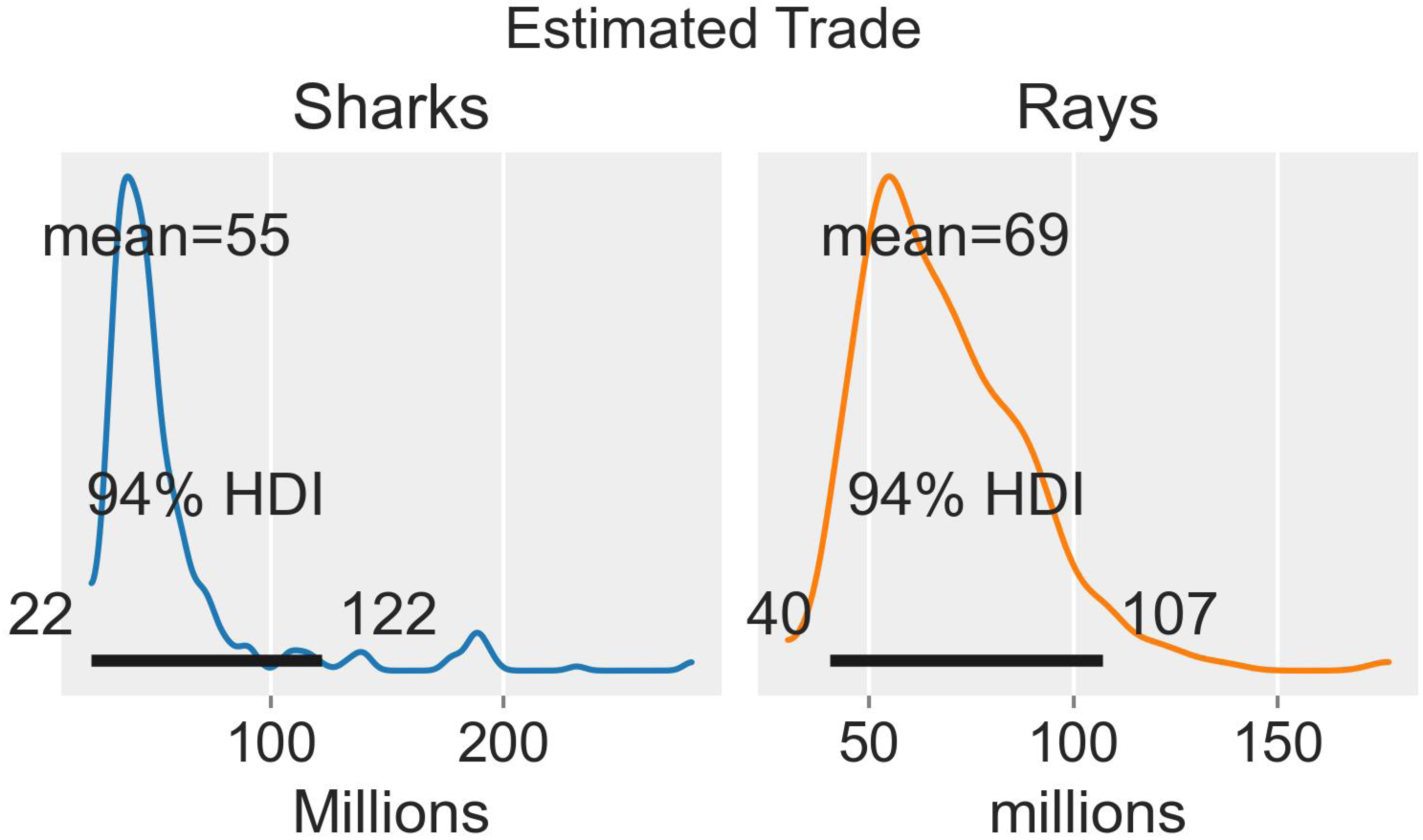
Estimated number of individual a) sharks and b) rays within the global meat trade annually. These conservative estimates stem from dividing model-estimated biomass of species in trade by their average mass at 75% maturity (from fishbase.org).

**Supplemental Figure 6.**
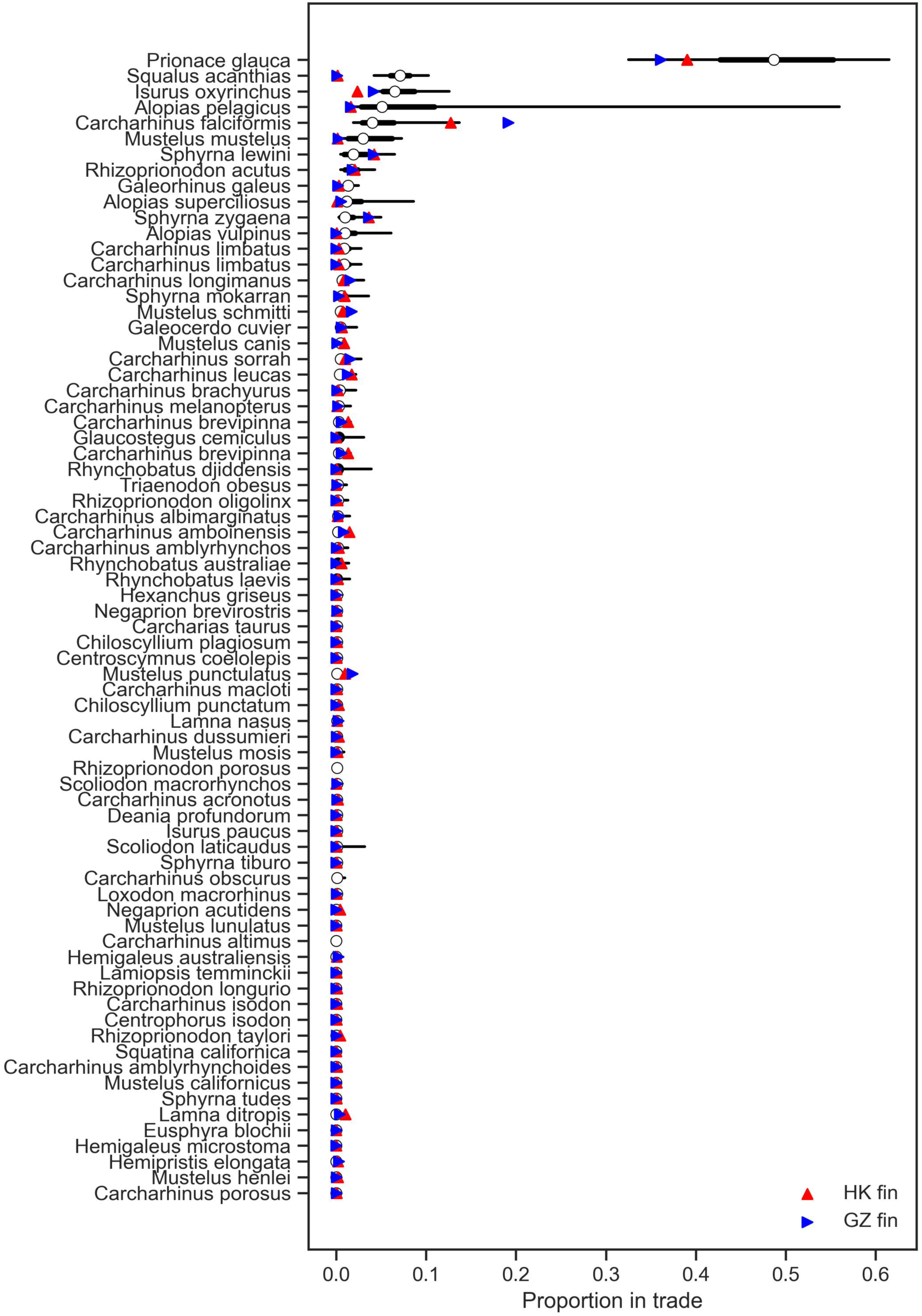
Comparison of species proportions for the top species estimated to be within the global meat trade (black) with proportions of species in the Hong Kong (HK fin, red up triangle) and Guangzu (GZ fin, right blue triangle) fin trades (*24*).

## Appendix A – Augmented elasmobranch landings

Some countries catching substantial numbers of elasmobranchs report only a small quantity to the FAO capture production database in elasmobranch categories (*23*). For example, Vietnam reported no landings in any elasmobranch category from 2012 to 2019, yet local published reports record landings approaching 12,000 tons/yr, in 2015 and 2016, from two separate provinces, comprising 0.2 to 0.5% of total national seafood landings (*41*). While this subset of countries does not report many elasmobranchs, reporting of miscellaneous marine fish with the three-letter code MZZ (Marine fishes NEI) provides information as to the scale of their likely elasmobranch landings, if an average proportion of total landings can be estimated. Therefore, to overcome the aggregation of elasmobranchs into the Marine fishes NEI category we developed a missing data imputation model to estimate the total latent elasmobranch landings from unidentified marine fish landings among all exporters in our analysis.

Landings data were taken from FAO Global Capture Production dataset for the period of 2012 to 2019 (*23*). Catches were classified as “elasmobranch” if they were in any taxonomic category associated with elasmobranchs, excluding chimaeras, and “finfish” if they were in the landings groups identified as “Flounders, halibuts, soles”, “Herrings, sardines, anchovies”, “Miscellaneous coastal fishes”, “Miscellaneous demersal fishes”, “Miscellaneous pelagic fishes” or “Tunas, bonitos, billfishes”. Landings in group MZZ were classified as “unknown”. All other landings were excluded, including invertebrates and any landings from freshwater regions and groups.

Total latent elasmobranch landings per country and year were modeled using a linear model. Mean log elasmobranch catch in each year and country was predicted as:

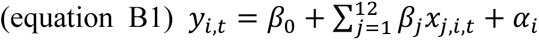

Where *y* is log(total shark catch + 0.01) in country *i* in year *t*; 𝛽_’_ is an intercept term, and 𝛽_)_ to 𝛽_)*_ are coefficients associated with *x*_1_ to *x*_12_, the predictor variables: (1) log(total finfish catch in country *i* and year *t* + 0.01); (2) logit(fraction of catch that was unknown across all years in country *i*); (3) the fraction of identified finfish catches that came from pelagic species in country *i*; (4) number of elasmobranch species occurring in the catch in country *i* according to the expert judgment priors; (5-7) the first, second, and third axes of a principle components analysis (PCA) of the proportions of fish catch from different gear types (industrial, artisanal, subsistence, recreational, bottom trawl, gillnet, hook and line, seine, trap, dredge, and other) from the Sea Around Us Project data (*42*); (8) the number of FAO areas fished by the country, and (9-12) a country continent group indicator (Africa, Americas, Asia, Europe, Oceania), and 𝛼_$_ a normally distributed country-specific random effect. Numerical variables were standardized by subtracting the mean and dividing by the standard deviation.

For combinations of country and year without unknowns, log elasmobranch landings (+0.01) were given a normal likelihood with a common residual standard deviation 𝜎. For combinations of country and year in which any unknown landings were reported, the landings were imputed from the same likelihood, subject to the constraint that latent elasmobranch landings must be between the reported elasmobranch landings and an estimated upper bound. The upper bound was calculated as reported elasmobranch landings plus unknown landings times the estimated fraction of the unknown that could potentially be elasmobranchs *f*. This fraction was a single estimated parameter given a beta prior with a mean equal to the total elasmobranch landings divided by total landings of elasmobranchs plus fish across all the FAO landings excluding unknown (mean = 0.017), and a coefficient of variation of 0.1, corresponding to 𝑓∼𝑏𝑒𝑡𝑎(98.346, 5907.2). This strongly informative prior was used because it is known from both empirical data and trophic ecology that elasmobranch landings are a small fraction of total fish landings in all jurisdictions. The linear model coefficients, residual standard deviation, and standard deviation of the country random effect were given vague priors (𝛽_0’_∼𝑁(0,10), 𝛽*_j_*∼𝑁(0,1), 𝜎∼𝑁(1), 𝛼*_i_*∼𝑁(0, 𝜏), 𝜏∼𝐸(1)).

The model was implemented using Gibbs sampler in JAGS (*43*, *44*). This was done using the *dinterval*() likelihood function, which adds 0 to the log-likelihood if the log catch is in the interval and negative infinity otherwise. Code and datasets are available on GitHub (https://github.com/mamacneil/GSM_Science).

Among the countries that reported more than one million t per year of marine finfish, elasmobranch or unknown marine fish landings, the countries with a substantial fraction of unidentified catch were Vietnam, Thailand, Malaysia, Japan, Indonesia, India, and China (Figure B1a). Large shark catches were positively associated with finfish catches, PCA-2 from the gear types, which related to bottom trawls and gillnets, number of shark species caught, and fraction of fish catch unidentified with broad credible intervals on the effects of continent group (Figure B1b). Across all 63 countries, our model estimated 17% more elasmobranch landings than the reported landings, but these increases were concentrated in the seven countries with substantial unknown landings, especially Indonesia, China, and Vietnam (Figure B1c). This result was strongly driven by the informative prior on the fraction of the unidentified fish catches that could be elasmobranchs; an alternative model in which any proportion of unidentified catches could be elasmobranchs (not shown) found unrealistically high elasmobranch landings. There was a slight declining trend in the observed elasmobranch catches, which was also visible in the estimated values (Figure B1d).

**Figure B1.**
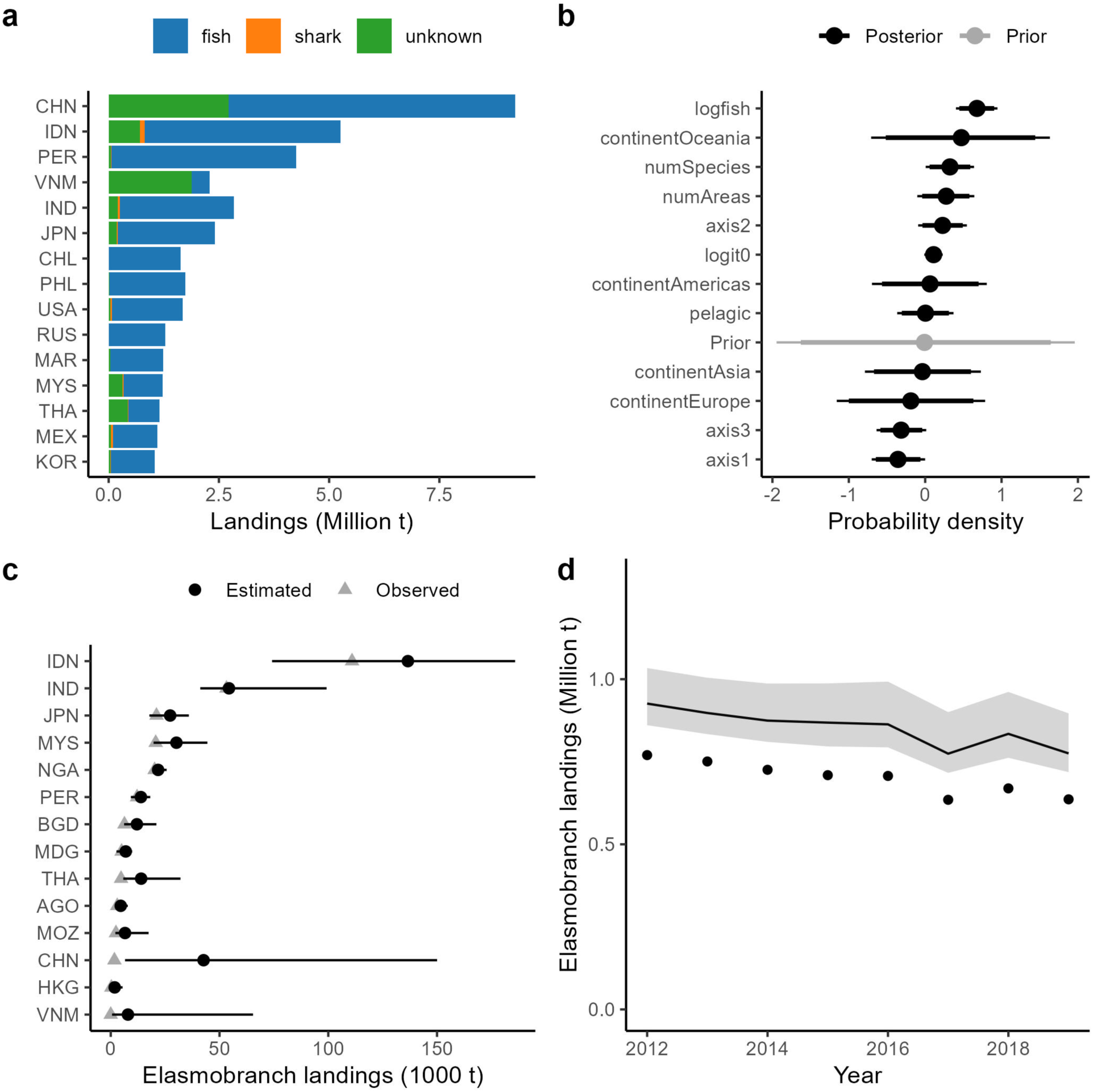
(a) Average landings per year in 2012-2019 by major fishing nations (>1 million t landings per year) of marine finfish (fish), elasmobranchs (shark), and the NEI category MZZ (unknown); (b) posterior densities of the regression coefficients (excluding intercept) from the log latent elasmobranch model, along with the prior (grey), showing 90% and 95% posterior density intervals; (c) median and 95% posterior density interval of annual mean elasmobranch catch for countries with substantial unknown landings, along with the reported catch (grey dot); and (d); the total across all 63 countries of estimated (95% posterior density interval) and observed elasmobranch catches (points).

## Appendix B – Statistical model structure

We chose to probabilistically decompose both landings and trade using a Bayesian statistical model (Supplemental Figure B1) that matched how data are reported. Because of the computational intensity required to estimate what amounts to many high-dimensional matrices, we chose to decompose landings first, using the posterior estimated latent landings as priors in our trade model, which is equivalent to running both sub-models simultaneously. PyMC code for the landings and models, along with all data handling, modelling, and graphics code can be found here: https://github.com/mamacneil/GSM_Science

The general model design is provided in Supplemental Figure B1 and the variables are defined in Supplemental Table B1. The notation for our model is as follows – with **data in bold**, for exporter (*e),* importer (*i*), species (*s*), taxon (*t*), year (*y*), shark or ray (*k*),

*Trade model*

The likelihood for the log trade of sharks or rays (*k*) from exporter *e* to importer *i* in year *y* is normal:

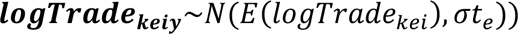

where the log-mean is the log of the sum over species of the expected exports from *e* to *i* of species that could be in *k*:

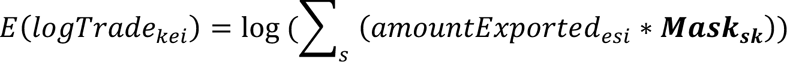

where the mask is the matrix of zeros and one that is used to remove species that should not be included. The log scale residual standard deviation is calculated as a linear regression from the Comtrade unreliability score:

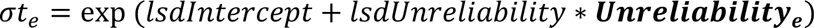

with priors for the coefficients of:

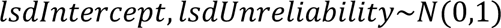

The quantity of each species exported from *e* to *i* is calculated as the estimated latent landings of the species by the exporter multiplied by the proportion of the species that is exported to *i* rather than consumed domestically or exported to another country:

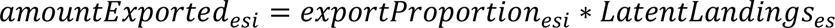

The export proportions are calculated from a linear model that includes the affinity of the importer for the species, a parameter specific to each dyad, and the strength of seafood trade from *e* to *i*, with non-trading dyads removed with a mask:

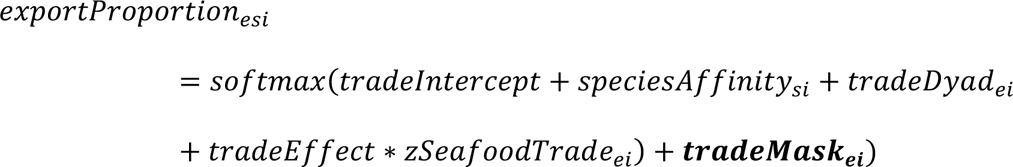

The coefficients have priors that are normal for the intercept and slope and zero sum normal for the categorical variables.

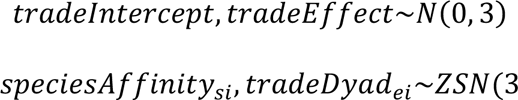

*Landings model*

The likelihood for landings identified to species is simply:

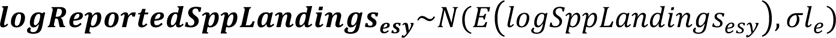

while landings estimated to higher taxonomic levels (*t*) have likelihood:

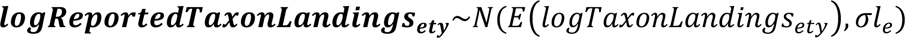

with a common log-standard deviation for all taxa within each exporter calculated as a log-linear regression from the BACI unreliability score:

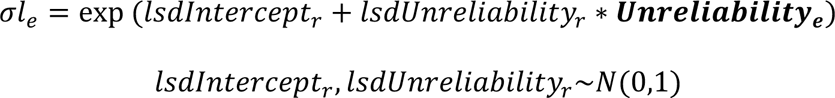

The expected log landings (identified to species) of species s in e in each year is the expected average annual landings adjusted by a ZSN error term:

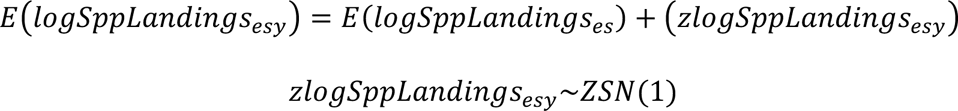

ZSN adds variation for each year that sums to zero over all years, akin to a random effect. The average annual landings are the latent landings multiplied by the proportion identified to species:

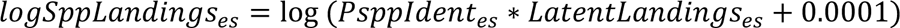

were proportion identified to species is estimated as a logit scale parameter for each species and exporter excluding non-existent species with a mask.

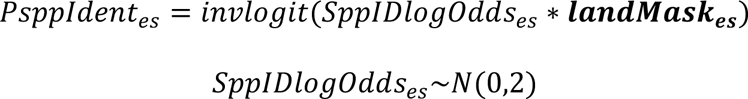

The log expected landings for each taxonomic group (above species), is similarly calculated as the average annual taxon landings plus a ZSN error term:

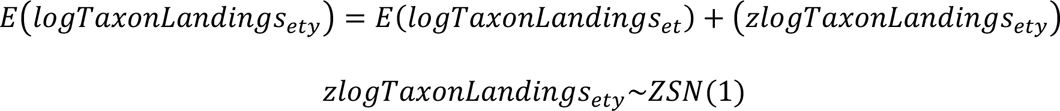

with the log taxon landings calculated as the sum over the species in the taxon of the landings of the species times the proportion of the aggregated (i.e. not identified to species) landings of species *s* reported in taxon *t*:

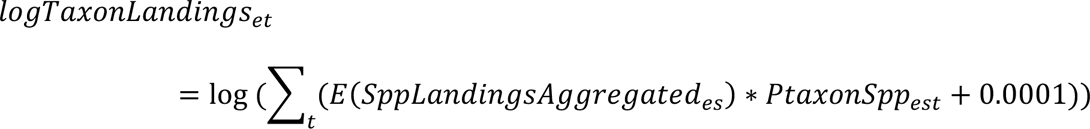

where the aggregated landings are just the fraction not identified to species:

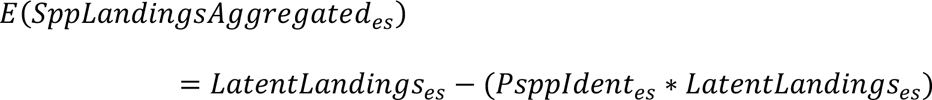

The proportion of aggregated landings of species *s* caught by exporter *e* and reported in taxon *t* has a different formula depending how many taxa the species could be allocated to in that exporter:

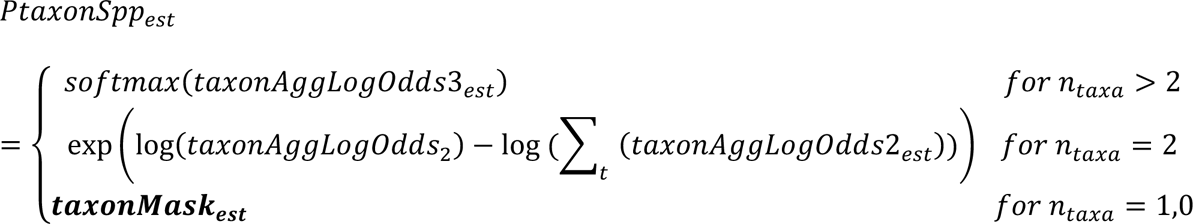

where, for two taxa, the log-odds of being allocated to one category verssus the other is just an estimated parameter:

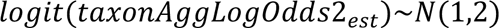

and for more than two categories the log-odds is an estimated parameter with a prior mean of 1 and magnitude adjusted using a linear model based on the number of taxa possible in the country (nTe) to account for the fact that larger numbers of categories influence the variance of the proportions in each category:

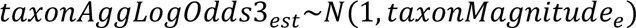

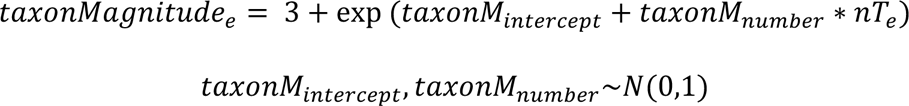

The latent landings of each species in each country is just the total landings times the species proportions:

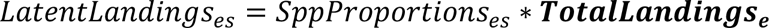

and the proportions are calculated from a softmax of the priors adjusted by a mask for species that exist in the exporter’s fishing areas:

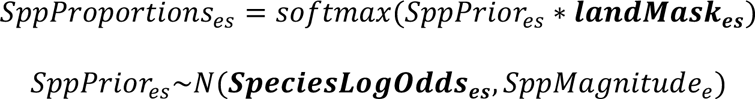

The prior for each species in each exporter has a mean from the logOdds elicited from experts, and a magnitude adjusted using a linear model based on the number of species possible in the country (nSe):

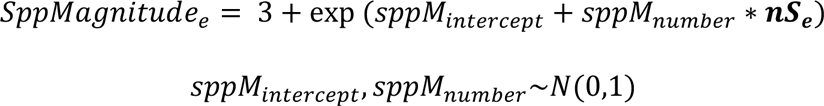

The likelihood for observed landings from the field studies associated with this project is:

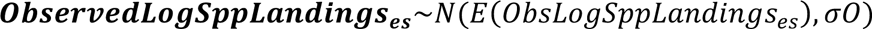

With an estimated log-standard deviation

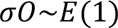

And log-mean calculated from the total landings in the country times species proportions.

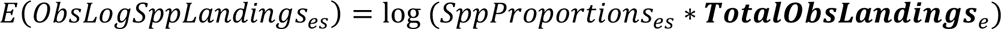

This notation matches the diagram in Supplemental Figure B1. The rationale in terms of our model structure is that only a limited number of species can comprise each level of taxonomic aggregation, from only a few species per genus and with an increasing number of additional species at higher taxonomic levels up to ‘elasmobranch’ (all species possible). The model recognizes this and explicitly integrates prior information for each species with corresponding permissible reported catch at each taxonomic level. In this way, species are allocated only to permissible taxonomic group aggregations.

One important consideration in all this is that the dimension of XX and YY are not constant among nations – each has a different species composition of varying dimension – which means that the prior variances for XX and YY are not constant. As such, out choice of prior variance is crucial – if we were to fix the variances it would suggest that we expect the log-odds to remain relatively stable and close to zero, regardless of the number of species. Instead, we allowed the prior variances to increase with the number of species present, meaning the logits can have larger magnitudes as needed, leading to higher log-odds in higher dimensions than the fixed variances would allow.

**Supplemental Figure B1.**
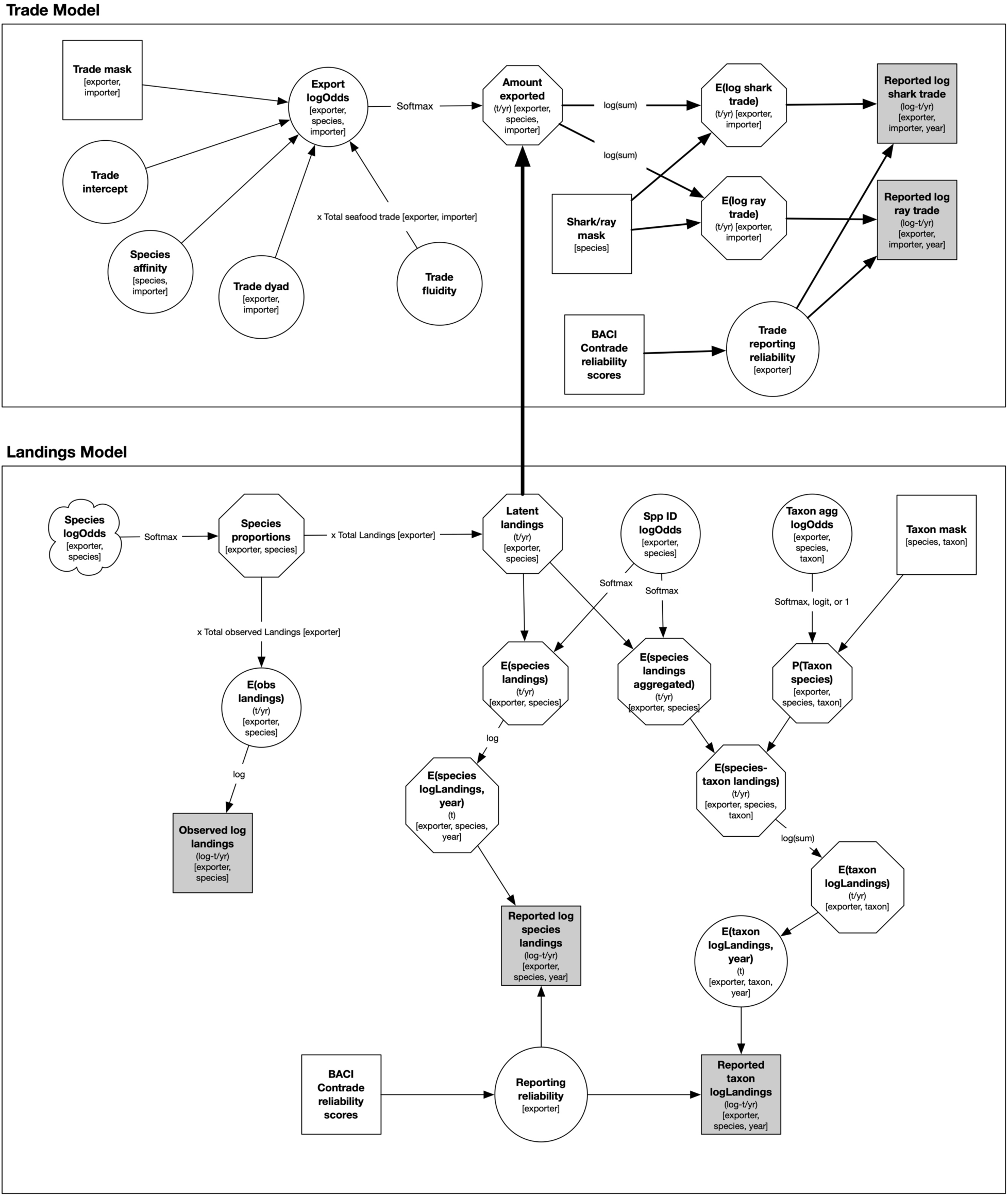
Modelling schematic of shark and ray meat landings and trade. Circles represent estimated quantities; squares represent data; cloud represents elicited priors; octagons are deterministic nodes; circles are stochastic nodes. Thick arrow indicates the use of latent landings posteriors from the landings model as priors used in the trade model, run sequentially due to computational constraints but mathematically equivalent.

**Supplemental Table B1.** Model variables used to quantify shark and ray meat landings and trade.

| Node | Type | Rationale | Units |
| --- | --- | --- | --- |
| *Landings model* |  |  |  |
| Species logOdds | Elicited information, stochastic node (prior) | Where landings composition is typically unreported, in-country experts were consulted about the relative abundance of landings by species, expressed on the log-Odds scale | Likert scale (*never, rarely, occasionally, some, common, a lot*) converted to logOdds (-4.5, -3.5, -2.5, -1, 1, 4). |
| Species proportions | Deterministic node | Proportions of each species in total landings per exporter | Probability (0-1) scale |
| Total landings | Data | Average total elasmobranch landings per exporter reported to FAO from 2012-2019 | tonnes per year |
| Latent landings | Deterministic node | Estimated average biomass of each species landed in each exporting country each year | tonnes per year |
| Spp ID logOdds | Stochastic node (prior) | log-Odds of each species in each country being reported to FAO at the species level | Probability (0-1) scale |
| Taxon agg logOdds | Stochastic node (prior) | log-Odds of each species in each country being aggregated up to each hierarchical taxon level | Probability (0-1) scale |
| Taxon mask | Data | A mask to exclude impossible species from each level of taxon aggregation | 0,1 |
| E(obs landings) | Deterministic node | Expected biomass of observed landings by species per exporter | tonnes |
| E(species landings) | Deterministic node | Expected biomass of landings reported by species per exporter | tonnes |
| E(species landings aggregated) | Deterministic node | Expected biomass of reported taxon aggregated landings by species per exporter | tonnes |
| P(Taxon species) | Deterministic node | Probability of each species in each country being aggregated up to each hierarchical taxon level | Probability (0-1) scale |
| Observed log landings | Data | Field and literature-based biomass per species per country | log(tonnes) |
| E(species logLandings, year) | Stochastic node | Zero-centred random effect for annual variation in reported log landings | log(tonnes) |
| E(species-taxon logLandings, year) | Deterministic node | Expected biomass of species contributing to reported taxon log-landings per exporter | log(tonnes) |
| E(taxon logLandings) | Deterministic node | Expected biomass of average reported taxon log-landings per exporter | log(tonnes) |
| Reported log species landings | Data | Biomass of species level reported landings per country per year | log(tonnes) |
| E(taxon logLandings, year) | Deterministic node | Expected biomass of reported taxon log-landings per exporter per year | log(tonnes) |
| BACI Comtrade reliability scores | Data | Relative reliability of reported trade data according to BACI; item 2.4 in https://www2.cepii.fr/PDF_PUB/wp/2010/wp2010-23.pdf | Standard deviation |
| Reporting intercept | Stochastic node (prior) | Intercept for linear model of landings reporting reliability | Standard deviation |
| Reporting slope | Stochastic node (prior) | Slope of relationship to BACI Comtrade reliability score for linear model of landings reporting reliability | Standard deviation |
| Reporting reliability | Deterministic node | Estimated standard deviation (uncertainty) around reported landings data | Standard deviation |
| Reported taxon log-Landings |  |  |  |
| *Trade model* |  |  |  |
| Trade mask | Data | A mask to exclude impossible trade between countries that do not trade shark or ray meat | 0,1 |
| Trade intercept | Stochastic node (prior) | Intercept for linear model of species trade | log-Odds |
| Species affinity | Stochastic node (prior) | Parameter to allow for species preferences among importers within linear model of species trade | log-Odds |
| Trade dyad | Stochastic node (prior) | Parameter to allow for differential trade between exporter and importer dyad pairs within linear model of species trade | log-Odds |
| Trade fluidity | Stochastic node (prior) | Parameter to allow for varying level of shark or ray trade relative to overall seafood trade within linear model of species trade | log-Odds |
| Export logOdds | Deterministic node | log-Odds of each species present in exporter landings being traded to each importer | log-Odds |
| Amount exported | Deterministic node | Biomass of annual trade for each species from exporters to importers | tonnes per year |
| E(log shark trade) | Deterministic node | Expected biomass of total trade in shark meat between exporter importer dyads | log(tonnes) |
| E(log ray trade) | Deterministic node | Expected biomass of total trade in ray meat between exporter importer dyads | log(tonnes) |
| Reported log shark trade | Data | BACI-corrected Comtrade data for sharks | log(tonnes) |
| Reported log ray trade | Data | BACI-corrected Comtrade data for rays | log(tonnes) |
| Shark/ray mask | Data | A mask to exclude rays from shark total trade and sharks from ray total trade | 0,1 |
| BACI Comtrade reliability scores | Data | Relative reliability of reported trade data according to BACI | Standard deviation |
| Trade reporting reliability | Stochastic node | Model-estimated standard deviation of trade date likelihood | Standard deviation |
| nSe | Data | Number of species reported per exporter | Number |
| nTe | Data | Number of taxon groups reported per exporter | Number |

